# Vascular K_ATP_ channel structural dynamics reveal regulatory mechanism by Mg-nucleotides

**DOI:** 10.1101/2021.05.15.444267

**Authors:** Min Woo Sung, Zhongying Yang, Bruce L. Patton, Barmak Mostofian, John Russo, Daniel M. Zuckerman, Show-Ling Shyng

## Abstract

Vascular tone is dependent on smooth muscle K_ATP_ channels comprising pore-forming Kir6.1 and regulatory SUR2B subunits, in which mutations cause Cantú syndrome. Unique among K_ATP_ isoforms, they lack spontaneous activity and require Mg-nucleotides for activation. Structural mechanisms underlying these properties are unknown. Here, we determined the first cryoEM structures of vascular K_ATP_ channels bound to inhibitory ATP and glibenclamide, which differ informatively from similarly determined pancreatic K_ATP_ channel isoform (Kir6.2/SUR1). Unlike SUR1, SUR2B subunits adopt distinct rotational “propeller” and “quatrefoil” geometries surrounding their Kir6.1 core. The previously unseen ED-rich linker connecting the two halves of the SUR-ABC core is observed in a quatrefoil-like conformation. MD simulations reveal MgADP-dependent dynamic tripartite interactions between this linker, SUR2B and Kir6.1. The structures captured implicate a progression of intermediate states between MgADP-free inactivated and MgADP-bound activated conformations wherein the ED-rich linker participates as mobile autoinhibitory domain, suggesting a conformational pathway toward K_ATP_ channel activation.

## INTRODUCTION

Dynamic regulation of K^+^ channel gating is a primary point of control for processes governed by electrical excitability. ATP-sensitive potassium (K_ATP_) channels, regulated by intracellular ATP to ADP ratios, transduce metabolic changes into electrical signals to govern many physiological processes^1^. They are uniquely evolved hetero-octameric complexes comprising four pore-forming inwardly rectifying potassium channel subunits, Kir6.x, and four regulatory sulfonylurea receptors, SURx, non-transporting members of the ABCC subfamily of ABC transporters^2^. Various Kir6.x/SURx combinations generate channel isoforms with distinct tissue distribution and function^3,4^. Best studied is Kir6.2/SUR1 channels, expressed in pancreatic β-cells, which control glucose-stimulated insulin secretion. Kir6.2/SUR2A channels are the predominant isoform in myocardium, while Kir6.1/SUR2B channels are the major isoform found in vascular smooth muscle. SUR2A and 2B are two splice variants of *ABCC9* that differ in their C-terminal 42 amino acids. In vascular smooth muscle, K_ATP_ activation leads to membrane hyperpolarization and vasodilation^5^, while inhibition or deletion causes membrane depolarization, vasoconstriction and hypertension^5–8^. Mutations in the vascular K_ATP_ channel genes (*KCNJ8* and *ABCC9*) cause Cantú syndrome^9–11^, a severe pleiotropic systemic hypotension disorder including hypertrichosis, osteochondrodysplasia, and cardiomegaly^12^.

K_ATP_ channel gating by intracellular ATP and ADP involves allosteric sites on both subunits. ATP binding to Kir6.x inhibits the channel. SURx, through induced dimerization of the paired nucleotide binding domains (NBDs), requiring MgADP bound to NBD2, and MgATP bound to non-catalytic NBD1, activates the channel^1,4,13^. Like all Kir channels, opening further requires PIP_2_ bound to Kir6.x^14–16^. Despite these commonalities, vascular Kir6.1/SUR2B K_ATP_ channels have distinct biophysical properties, nucleotide sensitivities, and pharmacology that differentiate them from other isoforms^17–19^. First, vascular K_ATP_ channel unitary conductance is half that of Kir6.2-containing channels. Second, vascular channels lack spontaneous activity, only opening in the presence of NBD-dimerizing Mg-dinucleotides/trinucleotides; in contrast, pancreatic or cardiac channels containing Kir6.2 open spontaneously in the absence of ATP. Third, once activated, vascular K_ATP_ channels are relatively insensitive to ATP inhibition, requiring mM concentrations to observe an effect, while their pancreatic or cardiac counterparts are blocked by ATP at μM concentrations. Lastly, the anti-diabetic sulfonylurea drug glibenclamide (Glib), which inhibits SUR1-containing pancreatic channels with high affinity, is ~10-fold less potent towards the vascular and cardiac channels containing SUR2. Glib has been shown to reverse defects from gain-of-function Cantú mutations in mice^20^. However, clinical application in Cantú patients is hindered by hypoglycemia from inhibition of pancreatic channels^21^. Structural mechanisms underlying unique biophysical, physiological and pharmacological properties among K_ATP_ channels are unknown.

Here, we report first cryo-EM structures for the vascular K_ATP_ channel, Kir6.1/SUR2B, in the presence of ATP and Glib. The structures show conformations not previously seen in pancreatic K_ATP_ channels prepared under the same condition^22–24^. First, unlike in Kir6.2, Kir6.1 cytoplasmic domains (CDs) were displaced from the membrane, too far to interact with PIP_2_ for channel opening. Second, unlike pancreatic channels, which have a uniform propeller-shaped conformation when bound to ATP and Glib^22,24^, vascular K_ATP_ channels held four distinct conformations, two resembling propellers and two quatrefoils, marked by varying degrees of rotation of SUR2B towards the core Kir6.1 tetramer. Importantly, a long segment of SUR not previously resolved in any K_ATP_ structures, linking NBD1 and transmembrane domain 2 (TMD2), was revealed within vascular K_ATP_ structures to mediate the cytosolic interface between SUR2B and Kir6.1. In particular, the linker’s unique 15-residue aspartate/glutamate (ED) domain^25^ established a nexus of interactions engaging SUR2B-NBD2 with Kir6.1-CTD. MD simulations showed MgADP binding to NBD2 was accompanied by substantial reconfiguration at this nexus, revealing the ED-domain provides a mobile autoinhibitory interaction that guards the transition of SUR2B from MgADP-free inactivated state to MgADP-bound activated state. Together our findings point to a structural pathway through which SUR regulates Kir6 channel gating.

## RESULTS AND DISCUSSION

### Structure determination of Kir6.1/SUR2B K_ATP_ channels with ATP and Glib

Vascular K_ATP_ channels were purified from COSm6 cells co-expressing rat Kir6.1 and SUR2B (97.6 and 97.2% sequence identity to human Kir6.1 and SUR2B, respectively). Channels were solubilized in digitonin, purified via an SUR2B epitope-tag, and imaged in the presence of 1 mM ATP (no Mg^2+^) and 10 μM Glib on graphene oxide (GO) coated grids, as described in Materials and Methods.

In vascular K_ATP_ channel structures as in pancreatic channels, we found SUR2B anchored to Kir6.1 via interactions mediated by TM1 of SUR2B-TMD0, and Kir6.1-TM1 (Fig.1). However, conformational deviations from 4-fold symmetry of the SUR2B were noted in 2D class averages (Fig.S1). To obtain clear SUR2B maps, we implemented symmetry expansion and extensive focused 3D classification of Kir6.1 tetramer with individual SUR2B (see Materials and Methods; Fig.S2), which isolated four 3D classes having identical Kir6.1 tetramer structures but different SUR2B orientations (Figs.1, S2). When symmetrized, two of the 3D classes, designated P1 and P2, resembled the pancreatic channel propeller conformations previously reported^22,24^. The other two, designated Q1 and Q2, resembled the “particular quatrefoil conformation” reported for human pancreatic K_ATP_ in which the SUR1 NBDs are dimerized^26^. Further refinement yielded cryoEM maps with overall resolutions of 3.4, 4.2, 4.0, and 4.2 Å for the P1, P2, Q1, and Q2 conformations, respectively (Fig.S3). The maps were sufficient to build a full atomic model for all of Kir6.1 minus the disordered C-terminus (368-424), with clear side-chain densities for most residues (see Fig.S3d), and also models for most of SUR2B (see Materials and methods for details). Densities for ATP, Glib, and some lipids were well resolved (Fig.S3d). Significantly, the Q1 conformation included definitive densities in SUR2B for L0, which is the linker connecting TMD0 and the ABC core, and also the N1-T2 linker, which connects NBD1 to TMD2; neither had been resolved in the human pancreatic K_ATP_ quatrefoil structure previously determined^26^. The P-like and Q-like conformations differ by a major rotation of the SUR2B-ABC core towards the Kir6.1 tetramer, clockwise when viewed from the extracellular side (Fig.1c,e). P1 and Q1 were the dominant particle populations within the P- and Q-like forms, respectively, differing from P2 and Q2 by degree of rotation and specific features. We first focus on structural differences between P1 and Q1, which provided the highest resolutions.

**Figure 1.**
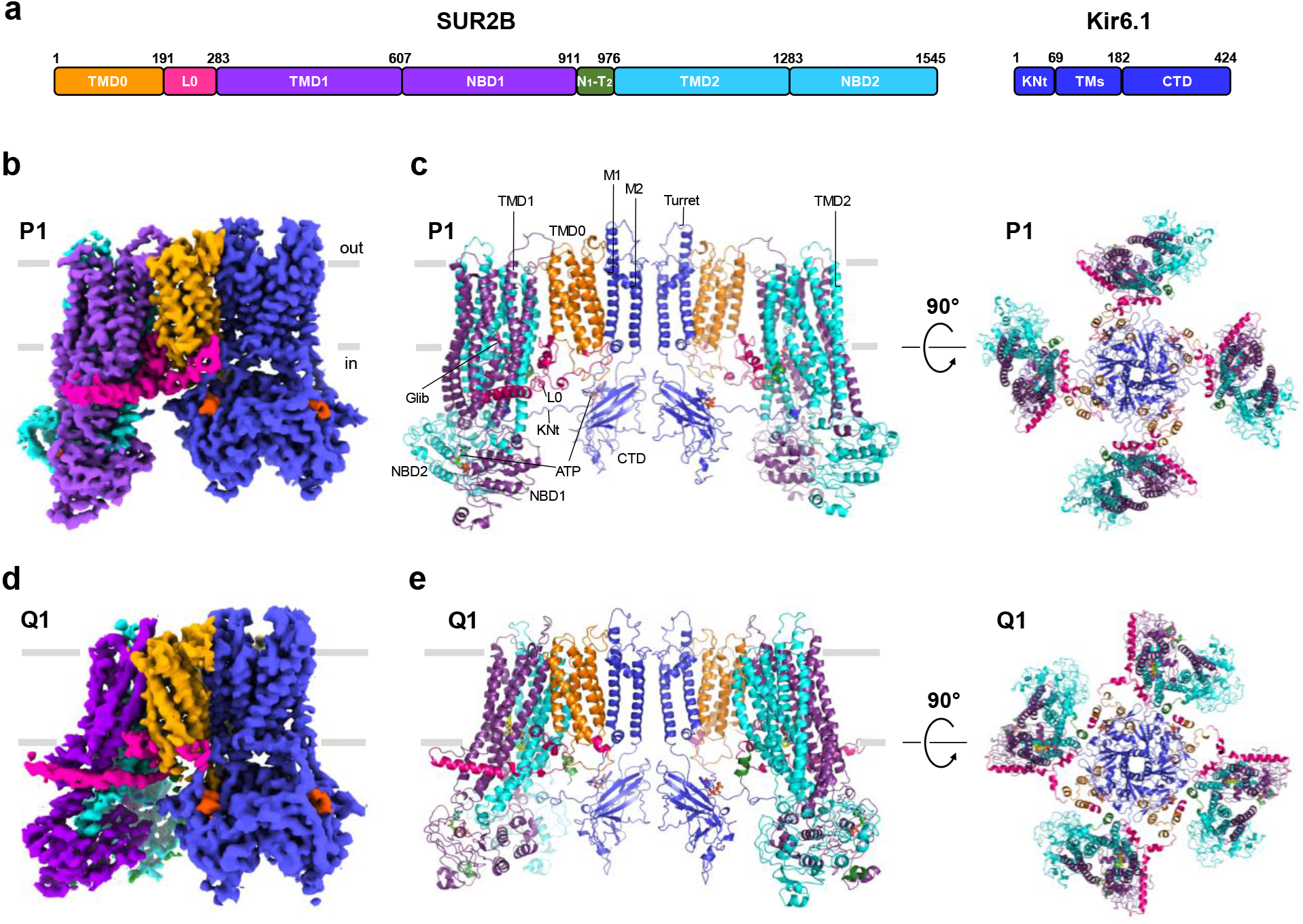
Structures of the vascular K_ATP_ channel in the presence of ATP and Glib. (a) Schematics of SUR2B and Kir6.1 domain organization. (b) CryoEM density map of (Kir6.1)_4_SUR2B P1, side view. (c) Four-fold symmetrized structure model of P1 viewed from the side (*left*) and the top (*right*). (d) CryoEM density map of (Kir6.1)_4_SUR2B Q1, side view. (e) Four-fold symmetrized structure model of Q1 viewed from the side (*left*) and the top, i.e. extracellular side (*right*).

### Structural correlates of Kir6.1 functional divergence

Although the Kir6.1 tetramer was similarly configured in all P and Q conformations for SUR2B, it included several features distinct from Kir6.2 in our published pancreatic channel structure determined under similar conditions with ATP and Glib (PDB: 6BAA). The Kir6.1 channel cytoplasmic domain (CD) was extended intracellularly away from the membrane by ~5.8 Å, and simultaneously counterclockwise-rotated (viewed from the extracellular face) by 12.4° (Fig.2A). The Kir6.x-CD is thus corkscrewed away from the membrane in Kir6.1/SUR2B, compared to Kir6.2/SUR1. Constrictions in the two cytoplasmic gates, namely the helix bundle crossing (F178) and the G-loop (G304, I305), indicate a closed Kir6.1 channel pore, similar to Kir6.2 under the same condition (Fig.S4). However, the distance between the helix bundle crossing gate and the G-loop gate is significantly larger in Kir6.1 due to the untethered CD.

**Figure 2.**
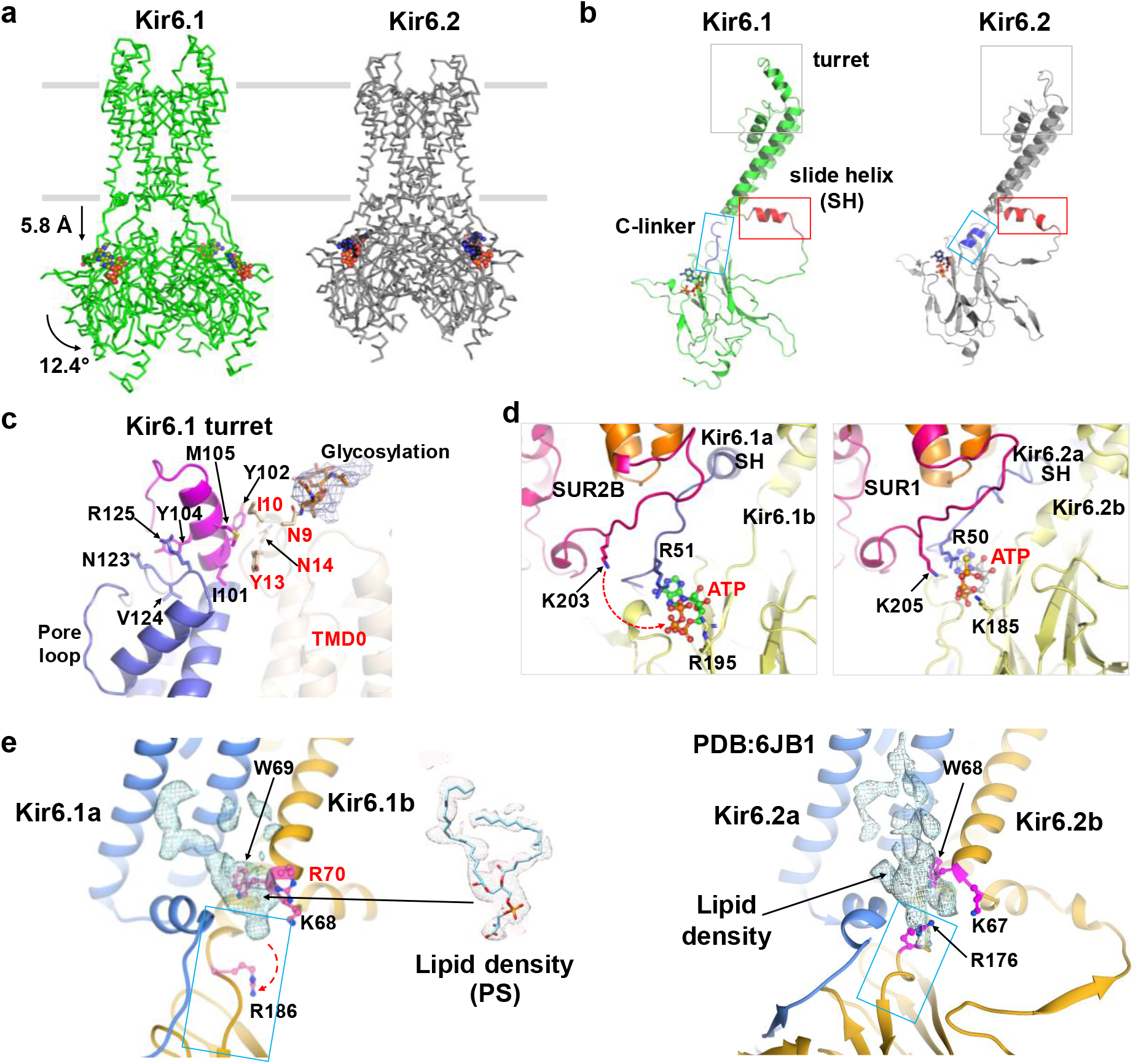
Structural comparison between Kir6.1 and Kir6.2. (a) Comparison of Kir6.1 and Kir6.2 showing translational and rotational differences in the CTD. (b) Major structural differences in the turret (grey box), slide helix (red box) and C-linker (cyan box) between Kir6.1 and Kir6.2. (c) Close-up view of the turret showing insertion of an additional 11 aa (magenta) in Kir6.1, which appears to be in position of interact with TMD0 of SUR2B (residues labeled in red). The density corresponding to glycosylation of N9 is fitted with two N-acetylglucosamines. (d) Close-up view of the Kir6.1 ATP binding site in comparison to Kir6.2 ATP binding site. R70 (P69 in Kir6.2) which could interact with negatively charged phospholipid is highlighted in red label. (e) Close-up view of the PIP_2_ binding site in Kir6.1 in comparison to that in Kir6.2.

In K^+^ channels, variations in the turret region surrounding the pore entryway have been shown to affect selectivity filter stability and ion conduction^27^. Compared to Kir6.2, the turret of Kir6.1 contains an extra 11 amino acids (^102^YAYMEKGITEK^112^). We found this sequence formed a helix and loop structure that extends the turret further out into the extracellular space (Fig.2b,c), potentially affecting conductance. Functional studies using Kir6.1-Kir6.2 chimeras previously identified residues in Kir6.1 thought to impart its smaller unitary conductance (Repunte EMBO J 1999). Among these, M148 in Kir6.1 (replacing Kir6.2-V138) is proposed to reduce pore entrance diameter; while N123 in Kir6.1 (replacing Kir6.2-S113) is hypothesized to impact an intersubunit salt bridge between R146 and E150, which in other Kir channels is formed by corresponding residues and critical for channel conduction^28,29^. However, our structure found M148 facing the pore helix (Fig.2c) rather than the entrance and that no significant difference exists in the adjacent pore diameters between Kir6.1 and Kir6.2, nor in their intersubunit salt bridges. Instead, we observed that N123-V124-R125 of Kir6.1, also implicated in the chimera studies, is uniquely located between the turret extension and the pore loop (Fig.2c), where it interacts with Y104 to potentially stabilize the turret and affect conductance.

We next assessed structural differences between Kir6.1 and Kir6.2 in two elements intimately associated with activity at ATP and PIP_2_ binding sites: the N-terminal amphipathic helix known as the slide helix (SH), and the connecting strand between TM2 and the C-terminal domain (CTD) called the C-linker (Fig.2b,d,e). In our Kir6.2 structure^22^, SH is bent halfway at the D58 position resembling a 3_10_ helix^30^. In contrast, SH in Kir6.1 formed a continuous helix extending toward the neighboring Kir6.1, thus compressing the PIP_2_ binding pocket. In Kir6.2, the C-linker forms a helix that tethers the CTD close to the membrane, which positions critical PIP_2_-binding residues such as R176 for PIP_2_ interaction. Rather different, the C-linker in Kir6.1 unwound into an unstructured loop stretching towards the cytoplasm, which deflected R186 (corresponding to Kir6.2-R176) away from the PIP_2_ binding site (Fig.2b,e). Without additional PIP_2_ in our sample, the PIP_2_ binding site nevertheless contained strong non-protein cryoEM density fitted by phosphatidylserine (PS), which may also bind in the PIP_2_ pocket, as recently shown in Kir2.1^31^. CryoEM densities matching ATP were also clearly resolved in Kir6.1 tetramers, at sites located between the N- and C-terminal domains of adjacent Kir6.1 subunits, matching sites in Kir6.2/SUR1 channels. However, ATP had fewer close residue interactions in Kir6.1 due to displacement of the Kir6.1-CD. In particular, in pancreatic channels SUR1-K205 (in L0) directly participates in binding ATP at its inhibitory site^23,26,32^, while the corresponding vascular channel residue SUR2B-K203 was displaced from potential ATP binding (Fig.2d). Thus, the constellation of ATP interactions was sparser and hence likely weaker when Kir6.1-CD was displaced from the membrane.

Taken together, translocation of the Kir6.1-CD away from the membrane compromised binding of both ATP and PIP_2_. This correlates well with the basal inactivity and reduced ATP sensitivity of the vascular K_ATP_ channel compared to Kir6.2-channels^33,34^. Rotation and downward movement of the Kir6.2-CD have been detected in minor subclasses of ATP- and Glib-bound pancreatic Kir6.2/SUR1 structures^23,24^, indicating similar dynamics occur but less stably persist. Moreover, translation and/or rotation of the CD is observed in Kir2, Kir3, and bacterial Kir channels ^35–38^, and recent cryoEM studies of Kir3 channels found that increased PIP_2_ concentrations shifts particle distributions towards those having CD tethered close to the PIP_2_ membrane sites^39^. Thus a common model of K_ATP_ channel activity involves channel opening dependent on PIP_2_ binding, which in turn depends on engagement by the Kir6.x-CD modulated by its vertical tranlocation/rotation. Accordingly, in vascular Kir6.1 channels, a greater energy barrier is involved in rotating the CD upward to interact with PIP_2_ than in pancreatic channels whose Kir6.2-CD is more stably tethered to the membrane. This explains why Kir6.2-containing pancreatic channels are spontaneously active, while Kir6.1-containing vascular channels are not. By extension, vascular channel activation by Mg-nucleotides likely involves SUR2B-controlled upward movement of the Kir6.1-CD (addressed below). Once activated by Mg-nucleotides, vascular K_ATP_ channels are highly stable, more resistant to PIP_2_ depletion than pancreatic channels^40^. In our Kir6.1 structure, the Kir6.1-R70 side chain directed toward the lipid density in the PIP_2_ binding pocket (Fig.2e) corresponds in Kir6.2 to proline (P69), a notable sequence variation that may contribute to the higher PIP_2_ affinity and stability of open vascular channels. Higher PIP_2_ affinity also accounts for long-standing results showing activated vascular K_ATP_ channels are much less sensitive to ATP inhibition, as increased PIP_2_ interaction reduces ATP inhibition in K_ATP_ channels^16^.

### SUR2B dynamics

Focused 3D classification resolved four distinct conformations, P1, P2, Q1 and Q2 showing variable SUR2B orientations (Fig.S5, video 1). P-conformations differed from Q-conformations by a large rotation of the ABC-core of SUR2B relative to the Kir6.1 tetramer (~41° between P1 and Q1, about the axis defined by N447 in TMD1 and N69 in TMD0, respectively; compared to 63° rotational difference between the propeller and quatrefoil conformations in human pancreatic NBDs dimerized channels measured from the equivalent residues). Within P and Q, P1 and Q1 particles predominated over P2 and Q2. Transitions from P1 to P2 and Q1 to Q2 involved alternative rotation stops: P1’s ABC-core was 10° further away from Kir6.1 than P2’s, while Q1’s ABC-core was 8° closer to Kir6.1 than in Q2. In short, Q1 was the tightest *quatrefoil*, and P1 the most extended *propeller*. 3D variability analysis in CryoSPARC (Fig.S6a) indicated SUR2B subunits moved independently between P-like and Q-like positions (video 2). Further multibody refinement in RELION3 revealed greater heterogeneity within Q1 conformations than in P1, indicating wider dynamic range (Fig.6b, video 3).

During rotation, the SUR2B-ABC core also tilts away from Kir6.1. Tilting elevated the ABC core TMD in the Q-conformations relative to P-conformations (by 2.6Å from P1 to Q1, measured at SUR2B-Y370; Fig.S5b). Between the pancreatic K_ATP_ *propeller* and *quatrefoil* forms (NBDs-dimerized), the entire ABC-TMDs elevate ~3Å without tilting^26^. Tilt in our Q-conformations may represent a partial transposition, to be completed upon NBD-dimerization. In the NBDs-dimerized pancreatic K_ATP_, quatrefoil is the dominant class. Here, Q-conformations were less common than P-conformations among vascular K_ATP_ channel structures in which the NBDs remain separated (Fig.S2, Table S1). Probabilities of SUR adopting P- or Q-like conformations therefore correlate with NBD dimerization state, although both occur regardless.

Rotation of SUR2B between P to Q conformations incorporated significant local structural changes. Extracellular contacts between TMB1 and TMD0 restructured both protein-protein and protein-lipid interactions (Fig.S7). Hydrophobic and electrostatic interactions in P1 are lost in Q1, including T338, L339 and F344 in the TM6-TM7 loop (TMB1), with L165 and R166 in TM5 (TMD0). Moreover, a phosphatidylethanolamine molecule moved from between TM2 and TM7 in P1, to between TM3 and TM16 in Q1, likely stabilizing TMD0 and TMB1 interactions. Also noteworthy, in the pancreatic channel structure SUR1 has an additional hydrophobic sequence (^340^FLGVYFV^346^), which anchors the TM6-TM7 loop to TMD0 (Fig.S7d)^23,32^. Absence of this sequence in SUR2B may impart flexibility that enables SUR2B to swing into Q-conformations not observed in SUR1 when ATP and Glib are bound.

### The SUR2B-L0 linker and the Glib binding pocket

Transition between P- and Q-conformations remodeled cytoplasmic structural elements including L0, the N1-T2 linker, and Kir6.1Nt, unexpectedly affecting interactions between SUR2B and Kir6.1. In SURx, L0 connects TMD0 to the ABC-core and is crucial to K_ATP_ gating^41–44^. In SUR2B, we obtained two distinct L0 conformations, corresponding to P and Q conformers. In SUR2B-P1, we observed continuous cryoEM density of L0 (Fig.3a). The well defined N-terminal portion lacked secondary structure. The central portion formed an amphipathic helix, inserted between TMD0 and TMB1. A C-terminal helix then extended along the periphery of TMB1, paralleling the membrane. In contrast, L0 of SUR2B-Q1 comprised a destabilized N-terminal portion in which aa 197-213 was unresolved; a central amphipathic helix shifted into the cytoplasm; and a C-terminal helix pulled away from the Kir6.1 core (Fig.3; video 4). In addition, lipids around the amphipathic L0 helix in P1-conformation were replaced by the descended amphipathic helix in Q1 (Fig.3a). Together, restructuring resulted in a marked decrease in contact area between SUR2B’s TMD0 (M1-R256, including lipids) and the adjacent ABC-core (A257-V1541 in P1; A257-A1543 in Q1), from 2106.2 Å^2^ in P1 to 1208.0 Å^2^ in Q1, which lowered the estimated free energy of formation at this interface from −43.0 kcal/M in P1 to −18.4 kcal/M in Q1 (calculated using PDBePISA^45^). As noted, the P1 conformer predominates among the ATP - and Glib-bound vascular K_ATP_ particles we observed. The simplest interpretation is that this interface is a principle determinant in maintaining SUR2B in P1 to a greater extent than Q1. The Q conformers would thus represent a divergent state in which a principal interface stabilizing a closed channel is compromised. It is worth noting that L0 in SUR1 (aa 192-262) is unresolved in the quatrefoil structure for the human pancreatic K_ATP_ channel, in which NBDs are dimerized (PDB 6C3O)^26^. Therefore the Q-like structures presented here may represent intermediary states that offer a glimpse into the conformational transitions of L0 that anticipate NBDs dimerization. The striking rearrangement of L0 likely results from the torque generated by rotation of the ABC core, and further permits the channel to undergo the conformational changes for gating.

**Figure 3.**
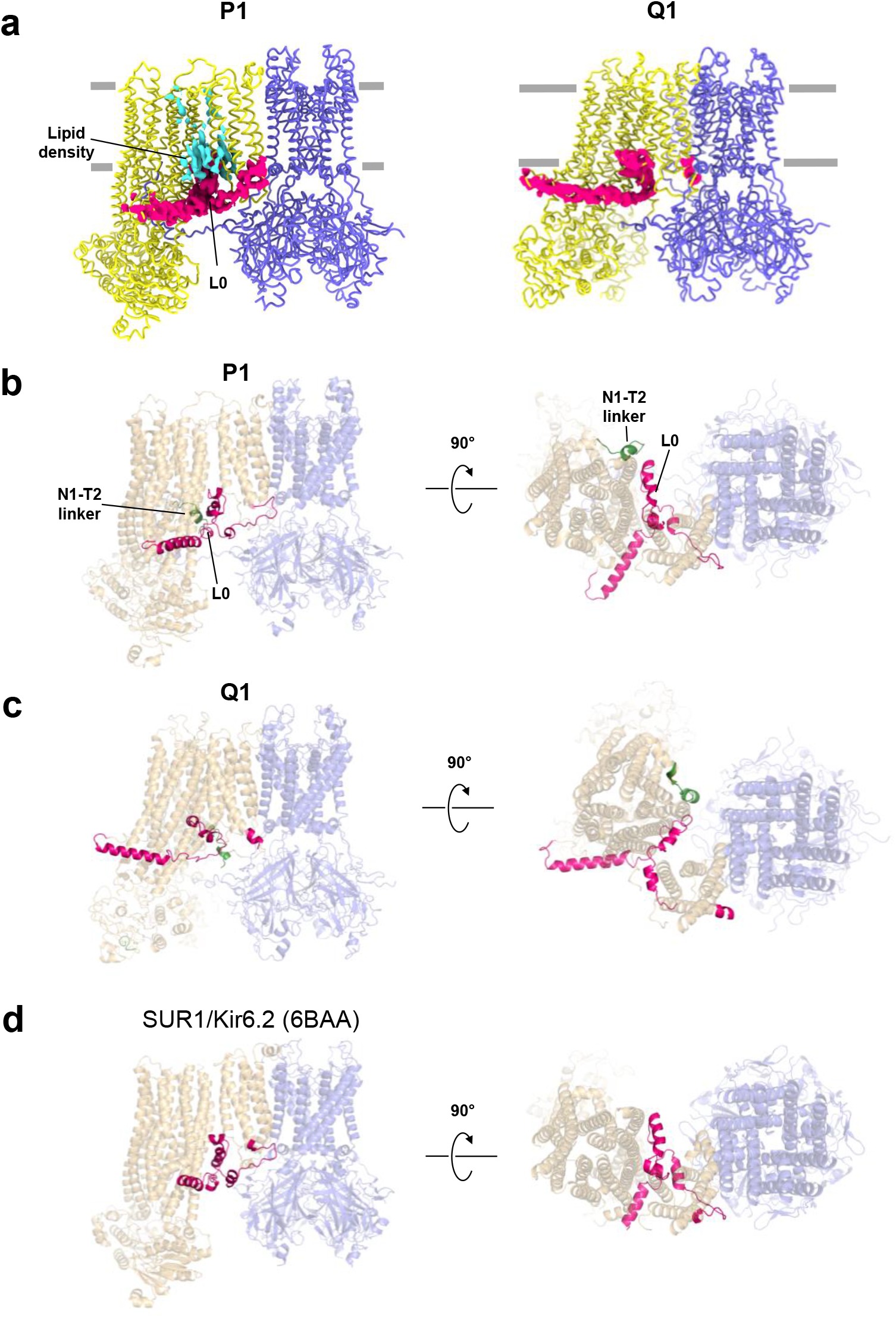
SUR2B-L0 undergoes structural remodeling from P1 to Q1 conformations. (a) Comparison of the L0 cryoEM density (hot pink) in P1 and Q1 conformations. Lipid density seen in P1 but absent in Q1 is shown in cyan. (b) Structure of (Kir6.1)_4_SUR2B in P1 conformations showing L0 (red) viewed from the side (left) and from the cytoplasmic side near the membrane (right). The N1-T2 linker visible in these views is shown in green. (c) Structure of (Kir6.1)_4_SUR2B in Q1 conformation viewed from the side and the bottom. (d) Structure of (Kir6.2)_4_SUR1 (PDB: 6BAA) bound to Glib and ATP for comparison.

Vascular K_ATP_ channels are inhibited by Glib but are ~10-50-fold less sensitive than pancreatic K_ATP_ channels^17,46^. Glib cryoEM density was well resolved in both P1- and Q1-conformations of the vascular K_ATP_ structure, where it bound within the same pocket of SUR2B (Fig.4) as in SUR1^24,47^. Also similar to pancreatic Kir6.1/SUR1 channels, cryoEM density of the distal KNt of Kir6.1 lay within the cleft between the two halves of the ABC core, and immediately adjacent the Glib binding pocket^47^. The structure model of the Glib binding site in P1 shows key interactions are largely conserved between SUR1 and SUR2B (Fig.4c). However, the binding pose of Glib in the Q1 conformation is compressed compared to that in P1, most particularly concerning the Y1205 side chain that was moved upward, which requires the 1-chloro-4-methoxy-benzene group to also move to avoid W423 in a neighboring helix (Fig4c). Also, an electrostatic interaction between chloride in Glib and nitrogen of R304 is eliminated in Q1. Worth noting, the SUR2B-Y1205 equivalent residue in SUR1 is S1238, and substitution of serine by tyrosine at this position has been shown to partly underlie Glib’s lower affinity inhibition of SUR2-channels^48^. Of particular interest, substitution of S1238 to Y in SUR1 renders the Glib inhibition of pancreatic Kir6.2/SUR1 channels, which is nearly irreversible, into a readily reversible inhibition similar to SUR2-containing channels^49,50^, suggesting the S1238Y mutation may affect Glib off rate. This may arise through steric hindrance from the flexible tyrosine side chain, as observed in the Q1-conformation.

**Figure 4.**
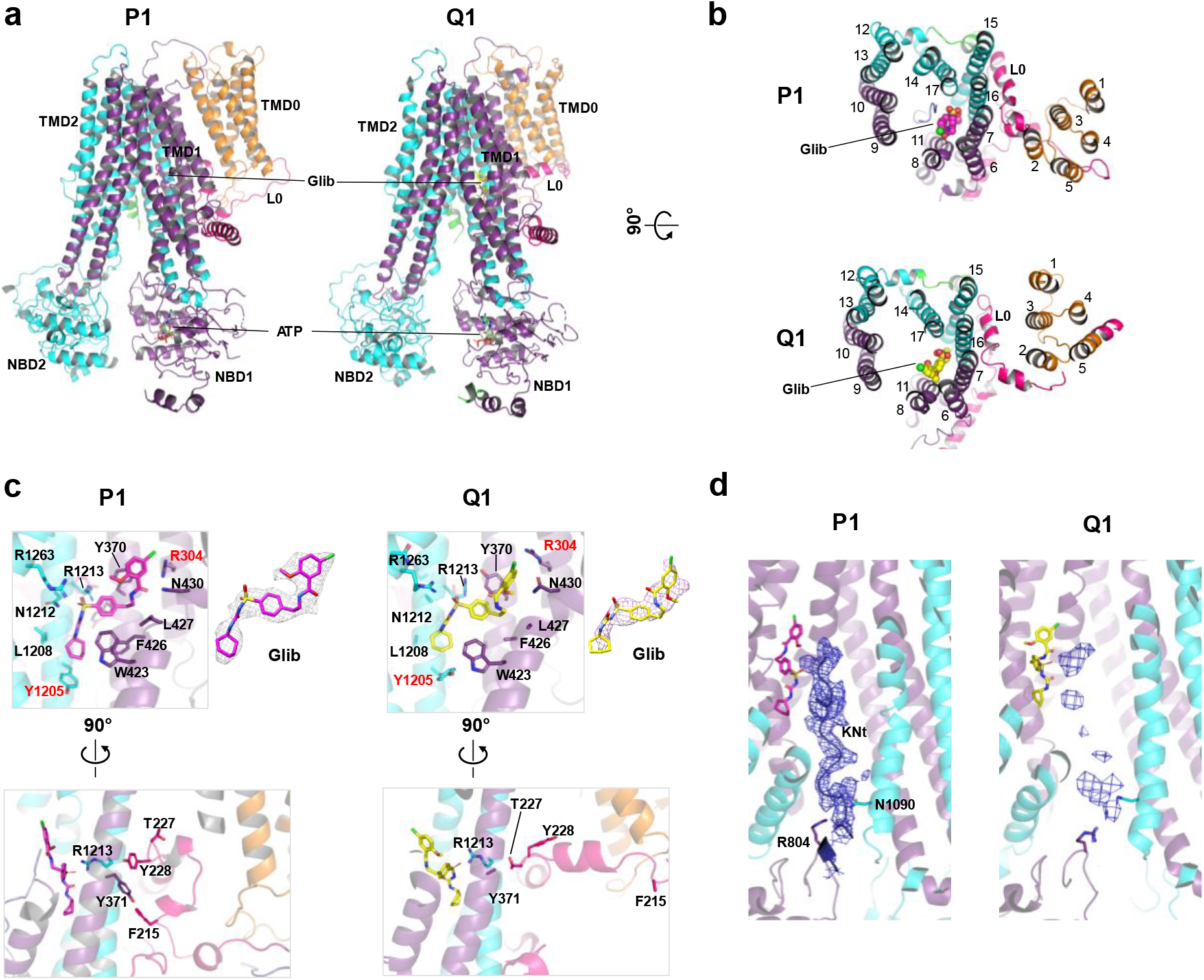
Comparison of the SUR2B Glib binding pocket in P1 and Q1 conformations. (a, b) Overview from the side and the top, respectively. (c) Close-up view of the glibenclamide binding site in P1 and Q1 conformations. Note the slightly different pose of glibenclamide. Two key residues different in SUR2B and SUR1 are highlighted in red (R304 and Y1205). CryoEM density with the glibenclamide structure model fitted into it is shown to the right of the binding site figure. Bottom: a different view of the glibenclamide binding site highlighting the changes in L0 residues that impact the glibenclamide binding site. (d) CryoEM density of the KNt in P1 and Q1 conformations. The KNt cryoEM density is stronger and allows modeling with a polyaniline chain. Note two residues in the NBD1 (R804) and TMD2 (N1030) sandwich the KNt to stabilize it in the central cavity between the two TMBs of SUR2B.

The P1 to Q1 translocation was accompanied by more substantial change to the opposite side of the Glib binding pocket, at F215, T227 and Y228 of the L0 linker (Fig.4c). Previous studies of L0 of SUR1 have shown that Glib binding indirectly involves Y230 and W232 (Y228 and W230 in SUR2B), which stabilize the TM helices lining the Glib binding pocket ^22^; mutation of these residues to alanine reduces sensitivity to Glib^49,50^. In SUR2B P1-conformation, we found the hydrophobic Y228 and W230 side chains, as well as F215 in the lower part of L0, similarly stabilized the TM helices along the Glib binding pocket (Fig.4c), as occurs in SUR1. Specifically, F215 lay buried in a hydrophobic cavity formed by W230 from L0 and Y371, F1207, L1206 from TMB1. However in Q1, L0 was significantly remodeled at the interface with TMB1. In particular, a loop segment including P218-Y228 seen in P1 is raised and transformed into a helix in Q1. This helical element newly filled the hydrophobic cavity between TMD0 and TMB1, otherwise occupied by lipids in P1 (see Fig.3a). As further consequence in Q1, Y228 and F215 in L0 are displaced from the cavity, and Y371 and T227 occupy the space vacated by the side chain of Y228. The movement of Y228 out of the cavity eliminates hydrophobic packing between L0 and the TM helices lining the Glib binding pocket, thus disrupting the integrity of the pocket in similar fashion to the Y230A mutational effect in SUR1 cited above^51^. Lastly, the density of Kir6.1Nt in the ABC core central cleft also differed between P1 and Q1 (Fig.4d). In P1, a strong continuous density of KNt was present, braced by R804 and N1090 of SUR2B, a pair of residues guarding entry to the cleft. The KNt density in Q1 was considerably weaker and discontinuous, indicating a more labile conformation that may contribute to weak Glib binding at its adjacent pocket^47,52^. In summary, as the SUR2B-ABC core changes from P-conformation to Q, L0 and Kir6.1Nt undergo remodeling that affects the Glib binding pocket.

### The NBD1-TMD2 (N1-T2) linker

In all published pancreatic K_ATP_ channel structures, the critical N1-T2 linker of SUR1 has remained unresolved^22–24,26,32,47,53^, suggesting dynamic instability. In the density map of our vascular Kir6.1/SUR2B channel from the P1 particle set, the C-terminal end of the N1-T2 linker was well resolved (Fig.5a) and yielded a polyalanine helical structure in the resulting model (residues 961-976). Density for the rest of N1-T2 (residues 911-960) remained largely unresolved in P1. However, the density for the entire linker was apparent in the map for our vascular K_ATP_ channel Q1 structure (Fig.5b), although resolution of residues 911-960 was insufficient for atomic scale modeling. Specifically, the linker extended from NBD1 through the space between the two NBDs, then continued through the gap between outer surface of NBD2 and the adjacent CTD of Kir6.1, before connecting to TMD2 (Fig.5c, Fig.S8). The location of the SUR2B N1-T2 linker contrasts sharply with locations of corresponding linkers in other ABCC proteins, including the Cl^-^ channel CFTR and the yeast cadmium transporter Ydf1p. In CFTR, the N1-T2 linker equivalent is known as the R domain, which is phosphorylated by PKA to allow CFTR gating by Mg-nucleotides. In unphosphorylated CFTR structure, the R-domain is wedged in the cleft between the two halves of the ABC core, preventing NBD dimerization^54^. In phosphorylated CFTR structure, the R-domain relocates to the outer surface of NBD1 (Fig.S8), which allows NBD dimerization, hence CFTR gating by Mg-nucleotides^55,56^. In the Yef1p structure, the N1-T2 linker is found at the outer surface of NBD1 similar to phosphorylated CFTR^57^, even though the NBDs are separate. The perculiar location of the SUR2B N1-T2 linker suggests the linker has adopted a separate role in regulating functional coupling between the SUR2B and Kir6.1.

**Figure 5.**
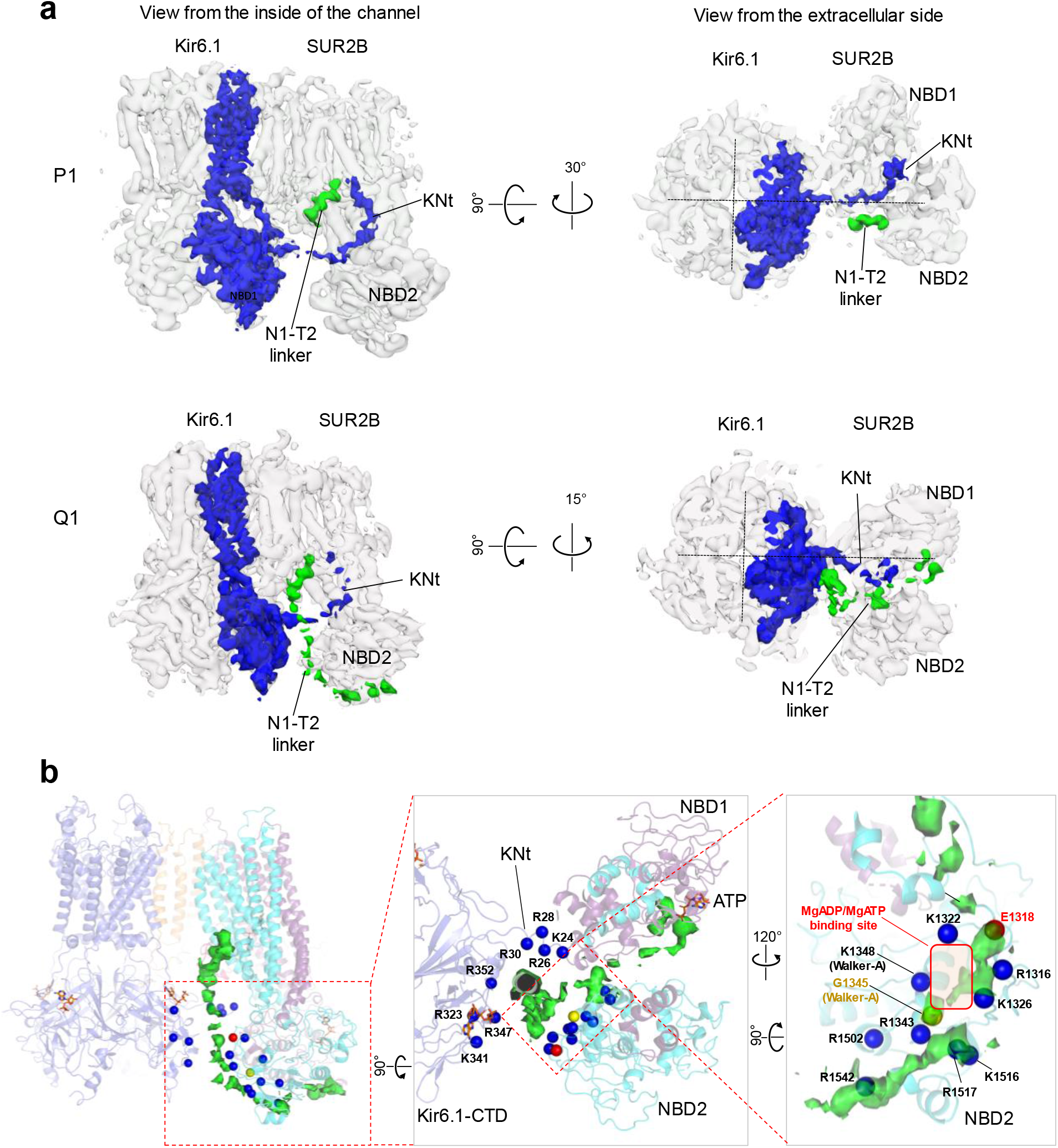
Comparison of cryoEM densities of Kir6.1 N-terminus and SUR2B N1-T2 linker in P1 and Q1 conformations. (a) Overall cryoEM density of (Kir6.1)_4_SUR2B in grey with density of one Kir6.1 and its N-terminus (KNt) highlighted in blue and density of the SUR2B N1-T2 linker highlighted in green. (b) Close-up view of the N1-T2 linker density in (Kir6.1)_4_SUR2B structure. Blue spheres are positively charged residues near the ED-domain. G1345 in the NBD2 Walker A motif and E1318 in the A-loop of NBD2 (^1315^VRYEN^1319^) are shown as reference points.

In SUR2, the N1-T2 linker includes at its C-terminal end a stretch of 15 amino acids consisting exclusively of negative charged glutamate and aspartate designated the ED-domain (947-961) (Fig.S8), which is unique among all ABCC proteins. Previous mutational studies have implicated the ED-domain in transducing MgADP binding in SUR2A to opening of Kir6.2^25^. Disruption of the ED-domain prevented the normal activation response to MgADP and to pinacidil, a potassium channel opener. In the Q1 structure, the density corresponding to the ED-domain is sandwiched between NBD2 and Kir6.1-CTD (Fig.5), and surrounded by positively charged residues from Kir6.1Nt, Kir6.1-CTD, and NBD2 of SUR2B (Fig.5b), an array of partners for electrostatic interactions. To understand the potential molecular interactions and their functional relevance, we employed MD simulations of the (Kir6.1)4-SUR2B Q1 structure.

### MD simulations reveal MgADP-dependent dynamic interactions between the ED domain, NBD2 and Kir6.1-CTD

To assess conformational dynamics of the ED domain and its interacting partners, and how they may be dependent on the nucleotide binding status at the two NBDs, we performed simulations under two conditions. In one, ATP is bound to Kir6.1 and NBD1 of SUR2B, as present in our cryoEM structure. In the second condition, Mg^2+^ is included with ATP bound at NBD1, and MgADP is docked into NBD2 (Fig.6a). In both conditions, Glib was omitted from the structure to allow the SUR2B TMDs to be free of constraint during simulations. To assess reliability, three independent 1μs simulations for each condition were carried out (Fig.S9a). In common with many biomolecular simulations, ours do not exhibit true equilibrium-like repeated fluctuations about mean values, although the combined 6 μs permitted structural inferences (Fig. 6c,e)^58^. The RMSF analyses showed high degrees of fluctuations of NBD1, the N1-T2 linker, and NBD2 (Fig.S9b), consistent with overall lower resolutions of these domains in cryoEM maps (Fig.S3b). However, particular interaction between the ED-domain and NBD2 depended on whether NBD2 was occupied by MgADP, and in turn those EDdomain-NBD2 interactions controlled direct interaction of NBD2 with Kir6.1-CTD.

**Figure 6.**
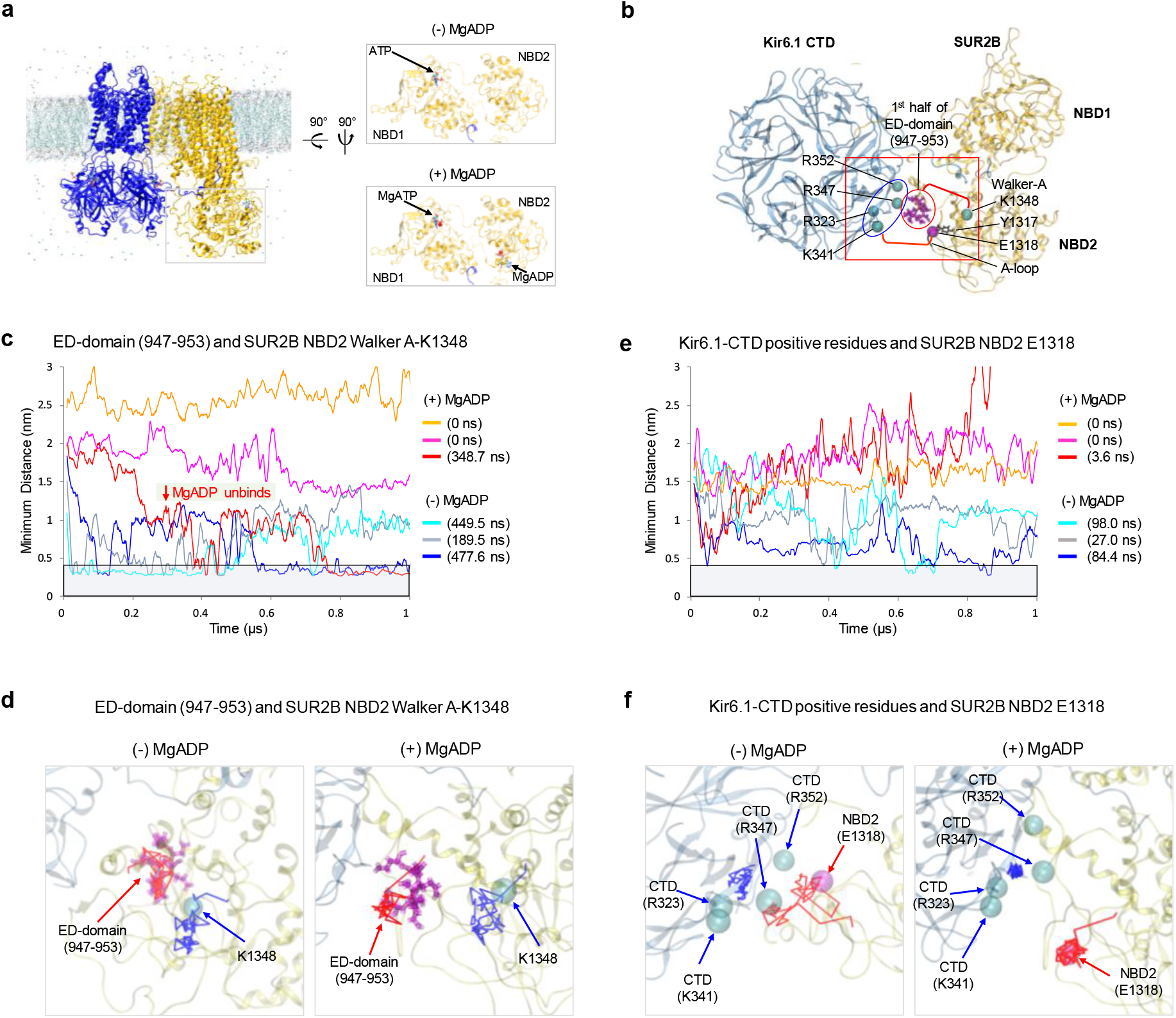
MD simulations of the ED domain dynamics in relation to SUR2B-NBD2 and Kir6.1-CTD. (a) MD simulation starting model (Q1) and conditions. In (-)MgADP condition, only ATP is present in NBD1. In (+)MgADP condition, MgADP is bound in NBD2 and MgATP is bound in NBD1. (b) Structural model marking residues of interest for distance analysis during MD simulations. These include the ED domain residues (magenta sticks in red oval) and the Walker A K1348 (cyan sphere) in SUR2B NBD2, and R323, K341, R347 and R369 (cyan spheres in blue oval) in Kir6.1 CTD and E1318 in SUR2B NBD2. The A-loop containing Y1317, which coordinates adenine ring binding of MgADP is also labeled. (c) Measurement of minimum distance between the side chain oxygen of any of the ED domain 947-953 glutamate/aspartate residues and the side chain nitrogen of K1348 in the three individual runs under both conditions. Note in one of the (+)MgADP runs (red) MgADP unbinds from NBD2 (marked by the red downward arrow). The grey bar marks the area where the distance is ≤ 4Å. The total dwell time in distance ≤ 4Å for each run is shown on the right. Note the plot was window-averaged with 10 ns scale and the dwell time was calculated with raw data which has 100 ps scale (the same applies to (e)). (d) Movement of the center of mass of the Cα of the ED domain residues 947-953 during simulation (red trace) relative to that of K1348 (blue trace). (e) Same as (c) except the distance measured is between side-chain nitrogen of Kir6.1-CTD residues R323, K341, R347, R352, and the side chain oxygen of E1318. (f) Same as (d) except the blue trace represents the center of mass of the Cα of Kir6.1 CTD residues R323, K341, R347, R352 and the red trace is the Cα of E1318.

During simulations, the ED-domain exchanged interactions between surrounding positively charged residues from Kir6.1Nt, Kir6.1-CTD, and NBD2 (see Fig.5b; videos 5,6). When MgADP is absent at NBD2, the first half of ED-domain (947-953) was most frequently in contact with SUR2B-NBD2 Walker A K1348; this was infrequent with MgADP bound at NBD2. To quantify a MgADP-dependence of the ED domain-Walker A interaction, we measured the minimum distances between the ED-domain residues 947-953 and K1348 throughout simulations, comparing results with or without MgADP at NBD2 (Fig.6b,c). In the absence of MgADP, side chain oxygens from ED residues were frequently within 4Å of the side chain nitrogen of K1348, supporting salt bridge or strong electrostatic interaction^59^. In contrast, in the presence of MgADP, ED residues remained too distant from K1348 for direct bonding. Moreover, in one MgADP simulation run in which MgADP dissociates (Fig.6c red trace ~300ns), the ED-domain subsequently moved to within 4Å of K1348, the distance frequently observed in simulations lacking MgADP (Fig.6c). The difference in ED-K1348 interactions between simulations is similarly evidenced by tracking the center of mass for Cα of ED residues 947-953 and the Cα of K1348 (Fig.6d).

Another outcome was that NBD2 frequently formed close contacts with Kir6.1-CTD in the absence of MgADP, but not when NBD2 included MgADP (videos 5,6). With no MgADP, a loop upstream of the Walker A motif in NBD2 (^1315^VRYEN^1319^, named A-loop for Aromatic residue interacting with the Adenine ring of ATP)^60^ frequently extended across the inter-subunit gap to interact with a cluster of positively charged residues in Kir6.1-CTD, including R323, K341, R347, R352 (video 5). In direct contrast, when MgADP bound to SUR2B-NBD2, the A-loop instead consistently interacted with MgADP at NBD2, far from the Kir6.1-CTD. The A-loop in SUR2B includes Y1317, which interacts with the adenine ring of bound MgADP at NBD2. Simultaneously, the dissociation of the ED-domain from Walker A K1348 that occurred with MgADP binding at NBD2 freed the ED-domain to move in between NBD2 and Kir6.1-CTD, There, the ED-domain interacts with positive-charged residues in Kir6.1-CTD that in the absence of MgADP interacted with NBD2 A-loop E1328 (videos 5,6). Effectively, the Kir6.1-CTD exchanges the A-loop for the ED domain, and stabilizes each conformation. Quantitatively, minimum distances measured between E1318 in A-loop and the four positive residues in Kir6.1-CTD documented the closer relation of A-loop and Kir6.1-CTD throughout the simulations in the absence of MgADP, than when MgADP was bound (Fig.6e). Minimum distance below 4Å sufficient for E1318 salt bridge formation was seen in all three runs lacking MgADP, but only transiently (3.6ns) in a single of the three runs with MgADP (Fig.6e). The nucleotide-dependent dynamics between the NBD2 A-loop and Kir6.1-CTD was also shown by tracking distance between the Cα of E1318 and center of the mass of the Cα for the Kir6.1-CTD positive residues (an example run for each condition shown in Fig. 6f).

The dynamic, tripartite interactions between the ED-domain, NBD2, and Kir6.1-CTD, and the dependence of these interactions on MgADP found in MD simulations significantly advances our understanding of the mechanism of SUR-mediated channel stimulation by Mg-nucleotides. In the absence of MgADP, the ED-domain has preferred interactions with NBD2 Walker A K1348, while the A-loop E1318 is engaged with Kir6.1-CTD. This hinders NBD2 from undergoing further conformational transition toward that of the NBDs-dimerized human pancreatic channel quatrefoil structure^26^, which shows SUR1-NBD2 further rotated towards NBD1 and also away from the positively charged residues in Kir6-CTD (PDB: 6C3O)^26^. Upon MgADP binding to NBD2, the ED-domain is dissociated from K1348 while the NBD2 A-loop becomes stabilized by the bound MgADP, unable to extend towards Kir6.1-CTD. As a sequence of results, the ED-domain is free to move towards other surrounding positively charged residues including those in Kir6.1-CTD, which further prevents the interactions between NBD2 and Kir6.1-CTD, thus allowing NBD2 to undergo further rotation towards dimerization with NBD1. Supporting this understanding, an ion pair formed by R347 in the Kir6.1-CTD, with E1318 in the A-loop of SUR2B-NBD2, has previously been reported to play a role in channel activation by MgADP and the potassium channel opener pinacidil^61^. Disruption of this ion pair by charge neutralization enhances MgADP/pinacidil gating, while charge swap restored wild-type like sensitivity to MgADP/pinacidil^61^. Our findings support the hypothesis that in order for NBDs to dimerize, interactions between SUR-NBD2 and Kir6-CTD must dissolve. Accordingly, disruption of the Kir6.1 R347-SUR2B E1318 salt bridge facilitates MgADP/pinacidil stimulation, as breaking the salt bridge promotes nucleotide binding at NBD2 and allows the further NBD2 movement needed for NBD dimerization and channel activation. The ED domain in particular, by interacting with Walker A K1348, acts essentially as a mobile autoinhibitory motif, akin to autoinhibition mechanisms in many kinases^62^, that occludes NBDs dimerization in the absence of MgADP and is deflected to permit dimerization when MgADP has bound to NBD2. In this way, the ED-domain functions as a gatekeeper to prevent unregulated channel activation in the absence of MgADP.

Taken together, our structures and MD simulations capture conformations that appear intermediate between the NBDs-separate inactive and NBDs-dimerized active states. Significantly, many of the Cantú-causing SUR2B mutations are of residues in TM12, including Y981 and G985 in the second elbow helix, and W1014, T1015, S1016 at the top (Fig.7). TM12 is connected to the N1-T2 linker (Fig.7d). Many other Cantú mutations are in domains interacting with TM12, including a series throughout TM13 (F1035, C1039, C1046, S1050, and M1056), as well as H1001 in TM12 and R112 in TM14, which interface TM13 (Fig.7c). One of the most frequently mutated residue R1150 of TM15 is adjacent to the structured helix portion of the N1-T2 linker, C-terminal to the ED-domain (Fig.7d). The interconnectivity of these residues and their association with the N1-T2 linker suggest they may in common govern a critical conformational change during channel gating by Mg-nucleotides at the NBDs.

**Figure 7.**
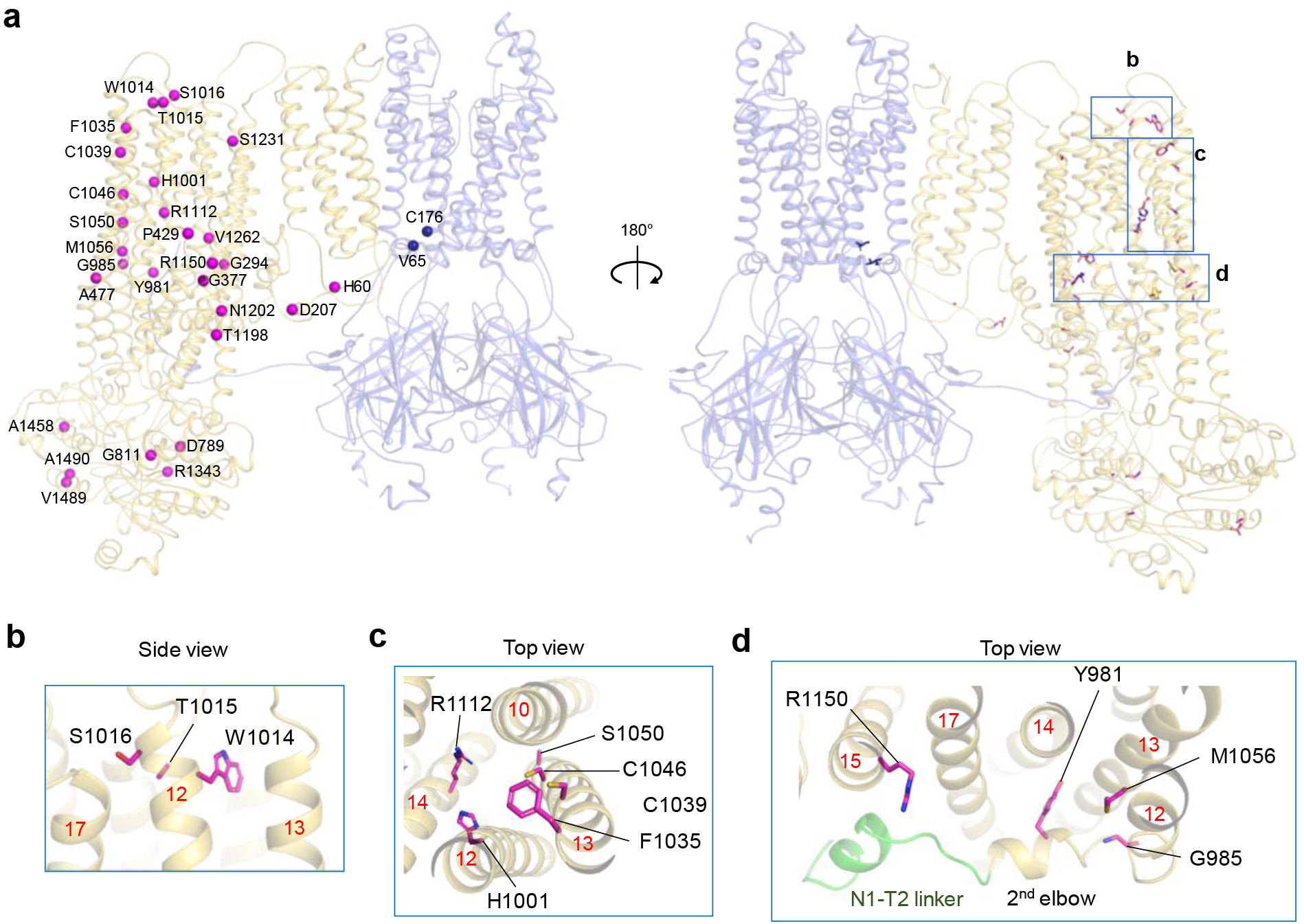
Residues mutated in Cantu patients mapped onto the Kir6.1/SUR2B channel structure. (a) Residues mutated are shown as blue (Kir6.1) or magenta spheres (SUR2B) in P1 conformation as spheres (left) or in stick model (right). Rat SUR2B numbering is used. Corresponding human mutations with rat residue in parentheses are as follows: H60Y (H60), D207E (D207), G294E (G294), G380C (G377), P432L (P429), A478V (A477), D793V (D789), G815A (G811), Y985S (Y891), G989E (G985), H1005L (H1001), W1018G (W1014), T1019E/K (T1015), S1020P (S1016), F1039S (F1035), S1054Y (S1050), C1043Y (C1039), C1050F (C1046), M1060I (M1056), R1116H/C/G (R1112), R1154G/Q/W (R1150), T1202M (T1198), N1206K (N1202), S1235F (S1231), V1266M (V1262), R1347C (R1343), A1462G (A1458), V1490E (V1489), A1494T (A1490). (b, c, d) Close-up side or top views of boxed regions labeled in the overall structure in panel a, right. In (d), the N1-T2 linker is colored green and labeled together with the second (2^nd^) elbow helix leading to TM12 of TMD2 in SUR2B. Red numbers mark the TM helices shown.

### Summary

Insights into how a particular complex operates is often gained by comparing related complexes anticipating that similarities and differences in structure and function will correlate. In this study we sought to determine the cryoEM structure of vascular K_ATP_ channels, composed of Kir6.1 and SUR2B, in the presence of ATP and Glib, for comparison to pancreatic K_ATP_ channel (Kir6.2/SUR1) structures determined with the same conditions. The novel structures we obtained reveal multiple elements showing distinct configurations that account for channel-specific conductances, ATP-inhibition, and drug sensitivities. In contrast, the serendipitous appearance of new quatrefoil-like conformations, and SURx linkers which have been missing in previous K_ATP_ structures and are now seen at critical domain interfaces, affords insights into the long-sought structural dynamics shared by K_ATP_ channels in regulating their activity. The Q-conformations adopted by SUR2B are most simply interpreted as transitional states between the inactive NBDs-separated and the active NBDs-dimerized SUR conformations.

The several conformations isolated from the cryoEM dataset, together with the dynamics revealed by 3D variability analyses and captured by MD simulations, suggest a model hypothesis for how Mg-nucleotide interactions with SUR2B activates Kir6.1 (Fig.8). In this model, individual SUR2B subunits transition between P- and Q conformations. In the Q-conformations and without Mg-nucleotides at NBD2, the ED-domain in the N1-T2 linker acts as an autoinhibitory motif that prevents unregulated activation. Specifically, ED-domain interaction with Walker A K1348 at NBD2 promotes electrostatic interaction between NBD2 A-loop and Kir6.1-CTD, which further corrupts the Mg-nucleotide binding site and also withholds NBD2 from dimerization with NBD1. Addition of Mg-nucleotides relieves autoinhibition imposed by the ED domain, coupling organization of the Mg-nucleotide binding site to liberation of NBD2 to rotate towards NBD1 for dimerization. Yet to be determined mechanisms are required to explain how dimerization of NBDs in SUR2B leads Kir6.1-CTD to move up to the membrane to interact with PIP_2_ for channel opening. The model would predict that inhibitory ligands such as Glib or stimulatory ligands such as Mg-nucleotides or the potassium channel opener pinacidil, are able to shift the equilibrium of SUR2B towards P- or Q-conformations to drive channel closure or opening, respectively. It is important to note that dimerization of the NBDs was not observed during the 1μs simulation in the presence of MgADP/MgATP; moreover, only one SUR2B is present in the simulations, which prevents consideration of potential structural impact of neighboring SUR2B subunits. Future structures with NBDs dimerized and MD simulations of the full channel are required to confirm and extend our understanding of K_ATP_ channel activation. This notwithstanding, we speculate the general scheme of the model applies to other K_ATP_ channels with variations to explain isoform-specific sensitivities for Mg-nucleotides and drugs. The structures presented here serve as a framework for understanding channel regulation and dysregulation, and will aid development of isoform specific pharmacological modulators to correct channel defects in Cantú and other diseases involving vascular K_ATP_ dysfunction.

**Figure 8.**
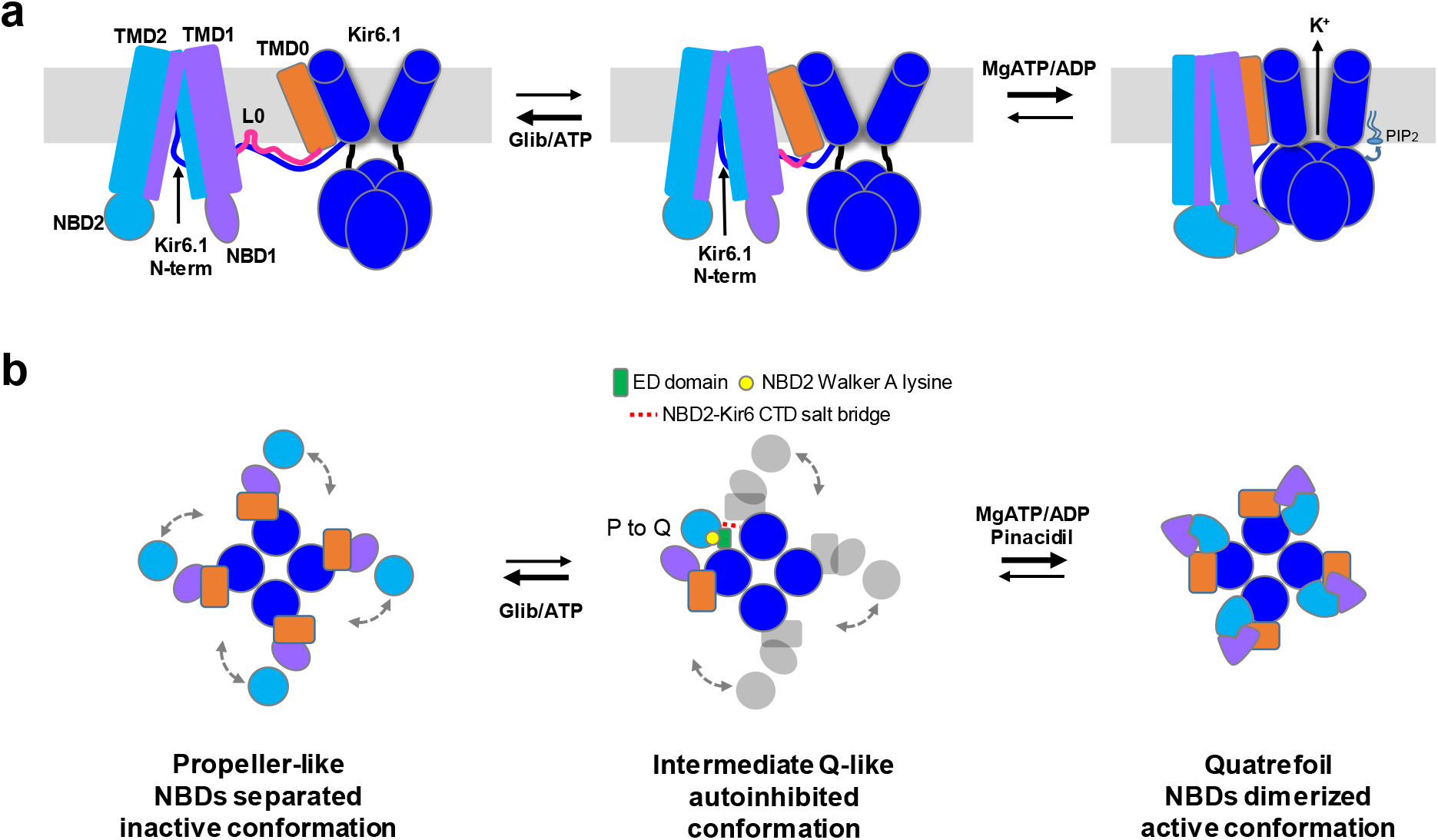
Proposed model of vascular K_ATP_ channel conformational dynamics. (a) Cartoon representation of channel side view and (b) top/down view in inactive P-conformation, Q-like intermediate conformation, and active, NBDs dimerized closed quatrefoil conformation. In the presence of Glib and ATP, the P-conformation dominates. Addition of MgATP/ADP promotes NBD dimerization, which is postulated to cause Kir6.1-CTD to move close to the membrane to interact with PIP_2_ for channel opening. In (b) individual SUR subunits undergo P-Q conformation transitions independently. In the absence of MgADP at NBD2, the ED-domain interacts with NBD2-Walker A lysine (1348). The A-loop E1318 in NBD2 forms salt bridges with positively charged residues in Kir6.1-CTD, preventing further rotation of NBD2 needed for NBDs dimerization, thus arresting SUR in an autoinhibited intermediate conformation. Increasing MgATP/ADP concentrations increases the probability of MgATP/ADP binding to all SUR2B subunits to release autoinhibition and promotes conformational change to the NBDs-dimerized quatrefoil state for channel activation.

## Materials and methods

### Protein expression and purification

Rat Kir6.1 and N-terminal FLAG-tagged (DYKDDDDK) SUR2B were first cloned into pShuttle vectors and then the AdEasy vector (Stratagene), and packaged into recombinant adenoviruses in HEK293 cells according to manufacturer’s instructions^63,64^. Note the pShuttle vector used for SUR2B contains a tetracycline-regulated promoter, necessitating co-infection of a tTA (tetracycline-controlled transactivator) adenovirus for SUR2B expression. COSm6 cells grown in 15 cm tissue culture plates were infected with the Kir6.1, SUR2B, and tTA adenoviruses using multiplicity of infections (MOIs) optimized empirically. Cells were cultured in the presence of 5 μM glibenclamide post-infection to stabilize the channel complex. At ~40-48 hours post-infection, cells were harvested by scraping and cell pellets were frozen in liquid nitrogen and stored at −80°C until purification.

For purification, cells were resuspended in hypotonic buffer (15mM KCl, 10mM HEPES, 0.25 mM DTT, pH 7.5) and lysed by Dounce homogenization. The total membrane fraction was prepared, and membranes were resuspended in buffer A (0.2M NaCl, 0.1M KCl, 0.05M HEPES, 0.25mM DTT, 4% trehalose, 1mM ATP, 5μM GBC, pH 7.5) and solubilized with 0.5% Digitonin. The soluble fraction was incubated with anti-FLAG M2 affinity agarose for 4 hours and eluted with buffer A (without trehalose) containing 0.05% digitonin and 0.25 mg/mL FLAG peptide. Purified channels were concentrated to ~250-300nM (222-267μg/ml) and used immediately for cryo grid preparation prior to cryo grid screening and data collection. The final sample applied to the grids contained 1mM ATP (no Mg^2+^) and 10μM Glib. The purity and quality of purified channels were assessed by SDS-PAGE and by negative stain of prepared grids.

### Sample preparation and data acquisition for cryo-EM analysis

To increase protein adsorption to the cryoEM grids, a graphene-oxide (GO) grid preparation protocol was developed by modification of a previously described method^65,66^. Briefly, gold quantifoil 2/1 grids were cleaned with acetone and glow-discharged for 60 seconds at 15 mA with a Pelco EasyGlow®, and 4μL of 1mg/mL Polyethylenimine (PEI) in 25mM HEPES pH 7.9 was applied to each grid and incubated for 2 minutes followed by washing with water. Then, 0.1 ~ 0.13 mg/ml GO was vortexed vigorously and applied to the grid and incubated for 2 minutes followed by two washes with water. The grids were used immediately for sample vitrification. Ice thickness was optimized by varying blotting time and also through extensive screening of the grid in order to find optimal regions. Two grids were imaged from the same purification and were prepared as follows: 3μL of purified K_ATP_ channel complex was loaded onto GO grids. The sample was blotted for 1s (blot force −10; 100% humidity) and cryo-plunged into liquid ethane cooled by liquid nitrogen using a Vitrobot Mark III (FEI).

Single-particle cryo-EM data was collected on a Titan Krios 300 kV cryo-electron microscope (ThermoFisher Scientific) in the Pacific Northwest CryoEM Center (PNCC) at Oregon Health & Science University, assisted by the automated acquisition program SerialEM. Images were recorded on the Gatan K3 Summit direct electron detector in super-resolution mode, post-GIF (20eV window), at the nominal magnification 53,000x (calibrated image pixel-size of 1.653Å; super-resolution pixel size 0.8265Å); defocus was varied between −1.3 and −2.8 μm across the dataset (Table S1). The dose rate was kept around 20 e^-^/Å^2^/sec, with a frame rate of 3 frames/sec, and 70 frames in each movie, which gave a total dose of approximately 40 e^-^/Å^2^. In total, 4130 movies were recorded.

### Image processing

Image processing was carried out similarly to that described previously^47^. The raw frame stacks were gain-normalized, aligned, and dose-compensated using Motioncor2^67^ with patch-based alignment (7 × 7) and without binning. CTF parameters were estimated from the aligned frame sums using CTFFIND4.1^68^. Particles were picked automatically using DoGPicker^69^ with a broad threshold range to reduce bias and high-pass filtered at 120 Å using *relion_image_handler–highpass* 120 command before 2D classification using RELION-3.0^70^. Classes with high signal/noise ratios displaying fully assembled complexes and those missing 1 or 2 SUR2B subunits were selected; the particles were re-extracted at 1.2554 Å/pix and 1.654 Å/pix for P and Q conformations respectively, then used as input for 3D classification in RELION-3.0 (see Fig.S2). Symmetry was not imposed at the initial step to include all potential conformations. Extensive 3D classification was performed to assess heterogeneity within the data.

### 3D variability using cryoSPARC

Motion corrected micrographs were imported to cryoSPARC and patch CTF estimation was performed. A total of 5,675,640 particles were automatically picked by template picker and the particles were subjected to 4 rounds of 2D classifications. Only classes that show full channel are selected and a total of 138,530 particles were used to generate an initial model, which was then subjected to homogenous refinement resulting in 5.5 Å. Then the refined particles were subjected to Non-uniform refinement and CTF Refinement which improved the resolution to 4.2 Å. To probe dynamic motions of the channel, 3D variability was performed and the result showed highly dynamic movements of SUR2B (Fig. S6a).

### Particle classification

Particles from whole channel 3D classification were refined with C4 symmetry imposed to best align the Kir6.1 pore. They were then subjected to C4 symmetry expansion to yield 4 fold more copies. Further refinement was performed without symmetry and mask to allow alignment of heterogeneous particles without any restrains. A mask corresponding to Kir6.1 tetramer and one SUR2B was generated using a published 6BAA model^22^ and Chimera to conduct focused 3D classification without particle alignment. This initial focused 3D classification yielded a total of 6 classes, with two most dominant ones (classes 1 and 6; Fig.S2) showing distinct SUR2B conformations (final 139,944 particles for P1 and 70,830 particles for P2). Class 3 also showed good map quality for the Kir6.1 pore with clear helical structure features; however, the density for the SUR2B subunit was smeary and weak. From 2D classification, it was apparent that the SUR2B domain has significant heterogeneity (Fig.S1c). Therefore further refinement of the whole channel was performed using particles pooled from class 2 to identify possible additional conformations. The focused refinement of class 2 was conducted with a generous mask to minimize restrains. Consistent with results from 2D classification, which showed quatrefoil-like conformation, the focused refinement revealed a quatrefoil-like conformation. To further probe heterogeneity in this subset of particles, a model of Kir6.2 tetramer and one SUR1 from previously published structure (PDB: 6BAA) were fitted into the quatrefoil-like map separately to generate a synthetic mask for the novel conformation using Chimera. Further 3D classification was performed against this subset of refined particles. From this, we identified two additional conformations (71,880 particles for Q1 and 22,038 particles for Q2). Further focused 3D classifications with higher regularization T values ranging between 20 to 30 were performed for propeller-like and quatrefoil-like conformations respectively. Focused refinement of the Kir6.1 tetramer plus one SUR2B subunit was carried out in RELION-3 against the whole channel for each of the four conformations after partial signal subtraction that removes signals outside the masked region and iterated focused refinement^71^.

Multi-body refinement was employed for all conformations to improve the map quality of the dynamic ABC-core domain and linker regions of SUR2B. Since 3D classification indicates significant motions of the ABC-module of SUR2B, we used a masking strategy that includes Body 1 (Kir6.1-tetramer (P25-D366) + TMD0 (M1-L197) of SUR2B) and Body 2 (ABC-core (Q213-A1548) of SUR2B plus KNtp (M1-K24) of Kir6.1), to probe dynamics of the SUR2B ABC core. Multibody refinement with this strategy was repeated with varying range of standard deviations of degrees on the rotations and pixels on the translations to rule out artificial motions. With the searching parameters in the rotations and translations of the two bodies used in the multibody refinement, the principal component analysis in the *relion flex_analyse* program revealed that approximately 17 and 27% of the variances in P1 and Q1 respectively are explained by the first eigenvector (Fig.S6b). Finally, multi-body refinement was followed by three to four rounds of CTF refinement and particle polishing steps in between refinements of all four conformations. Final maps were subjected to Map-modification implemented in *Phenix* ^72^with two independent half maps and corresponding mask and model as input. They were then sharpened with model-based auto sharpening with the corresponding model using *Phenix*, a step that was iterated during model building.

### Model building

In the reconstruction of the P1 conformation of the Kir6.1-tetramer and one SUR2B, there is good density for nearly every side chain of Kir6.1, TMD0, and the inner helices of the ABC core of SUR2B. To create initial structure model, TMD (32-171) and CTD 172-352) of Kir6.2 from 6BAA model, TMD0/L0 (1-284), TMD1 (285-614), NBD1 (615-928), NBD1-TMD2-linker (992-999), TMD2 (1000-1319) and NBD2 (1320-1582) of SUR1 from 6PZA model ^47^ were used for rigid body fitting of Kir6.1 and SUR2B respectively in cryo-EM map of P1 conformation. Then each domain was combined using Chimera and used as the template model. All side chains and missing loops were then built by SWISS-model^73^ based on the template according to the sequence of Kir6.1 and SUR2B. In the P1 density map, NBD1/2 and loop regions showed signs of disorder, thus most residues in these regions were modeled as polyalanines and manually refined into the map. Loops and linkers where the density is absent were removed from the model and the side chains of amino acids in flexible regions were also not modeled. In addition to protein density, two N-acetylglucosamine (NAG) molecules which are common cores of N-linked glycosylation are modeled in the distinctively large density at the side chain of SUR2B N9. Densities corresponding to lipids were also observed at the interfaces and were modeled with best fitting lipid molecules accordingly. The resulting model was further refined using *Coot* and *Phenix* iteratively until the statistics and fitting were satisfactory. The final model contains residues 1-366 for Kir6.1, and residues 1-1541 for SUR2B except two loop regions (619-665 and 733-740) in NBD1 and the linker between NBD1 and TMD2 (911-960).

To build the Q1 model, TMD and CTD of the Kir6.1 tetramer, and TMD0, TMD1-2, NBD1 and NBD2 of SUR2B models from P1 were fitted as separate rigid bodies into the map and then combined as a template model. Models of the helical bundles in the ABC-core of SUR2B showed significant deviation from the Q1 density map. The initial Kir6.1 tetramer plus one SUR2B model was therefore refined iteratively by manual fitting using *Coot* and *Phenix* with morph option to allow smooth distorting of the model for a better fit into the map. NAG molecules and lipids are also modeled as in the P1 model. TMD0 from a neighboring SUR2B is also modeled with L0 loop located below the membrane as in the P1 conformation. The final model contains residues 24-368 for Kir6.1, and residues 1-1543 for SUR2B except two loop regions (617-663 which corresponds to L1 linker and 733-740) in NBD1, NBD1-TMD2 linker (911-960) and a loop (1460-1462) in NBD2.

Initial models for P2 and Q2 were built by docking Kir6.1 tetramer, TMD0, TMD1-2, NBD1 and NBD2 from P1 and Q1 models into the cryoEM density maps. Side chains are removed from the template models due to lower map qualities and the models were further refined using *Coot* and *Phenix* iteratively until the statistics and fitting were satisfactory. Note Glib was not modeled in Q2 due to low resolution of the map or absence of the ligand in this conformation.

### Molecular dynamics simulations

All MD simulations were performed at all-atom resolution using AMBER 16^74^ with GPU acceleration. Initial coordinates were developed from the Q1 model (four Kir6.1 and one SUR2B) with one ATP bound at SUR2B-NBD1 and four ATP molecules bound to the Kir6.1 tetramer; Glib was removed so as to allow the TMDs to relax during simulations. Kir6.1 Nt (M1-K24) and N1-T2 linker (M910-D961) built using SWISS model ^73^ were fitted into the cryoEM density and refined with *Phenix*^12^. L1 linker (A616-E664) from the SWISS model was used without manipulation. Initial simulation of the protein model showed the NBDs deviating significantly from the starting model. Additional testing showed that including all the lipid molecules found in the Q1 model stabilized the interfacial interactions particularly between TMD0 and ABC-core. Therefore, lipids were included in the starting model for simulations.

The simulation starting structures were protonated by the H++ webserver (http://biophysics.cs.vt.edu/H++) at pH 7 and inserted in a bilayer membrane composed of 1-palmitoyl-2-oleoyl-phosphatidylcholine (POPC) lipids and surrounded by an aqueous solution of 0.15 M KCl. The optimal protein orientations in the membrane were obtained from the OPM database^75^. Both systems contain 657 POPC lipids and ~111,000 water molecules, resulting in a total of ~470,000 atoms. They were assembled using the CHARMM-GUI webserver^76–78^, which also generated all simulation input files.

The CHARMM36m protein^79^ and CHARMM36 lipid^80,81^ force field parameters were used with the TIP3P water model^82^. Langevin dynamics^83^ were applied to control the temperature at 300K with a damping coefficient of 1/ps. van der Waals (vdW) interactions were truncated via a force-based switching function with a switching distance of 10 Å and a cutoff distance of 12Å. Short-range Coulomb interactions were cut off at 12Å, long-range electrostatic interactions were calculated by the Particle-Mesh Ewald summation^84,85^. Bonds to hydrogen atoms were constrained using the SHAKE algorithm^86^.

The atomic coordinates were first minimized for 5000 steps using the steepest-descent and conjugated gradient algorithms, followed by a ~2 ns equilibration simulation phase, during which dihedral restraints on lipid and protein heavy atoms were gradually removed from 250 to 0 kcal/mol/Å^2^, the simulation time step was increased from 1 fs to 2 fs, and the simulation ensemble was switched from NVT to NPT. To keep the pressure at 1 bar, a semi-isotropic pressure coupling was applied that allows the z-axis to expand and contract independently from the x-y plane^87^. The simulations were then run for over 1 μs with a time step of 4 fs enabled by hydrogen mass repartitioning^88,89^.

### Analysis of pairwise distances

Pairwise distances from the side-chain oxygen atoms in residues 947-953 in the ED-domain to the side-chain nitrogen atom in residue 1348 of Walker-A, and from the side-chain nitrogen atoms in residues 323, 341, 347, and 352 of the CTD domain to the side-chain oxygen atom of residue 1318 in the NBD2 domain were measured from the simulated trajectories using the gmx pairdist tool in Gromacs 2019.4^90^.

The ED-domain to SUR2B NBD2 Walker-A and the Kir6.1-CTD to SUR2B NBD2 analyses (Fig. 6c, e) were performed by selecting at each time the residue pair with the smallest distance. This showed if any of the residue pairs were close enough to form a salt bridge, even if the particular pair involved in that bridge changed. In addition to minimum distances, individual distances were also analyzed (Fig. S9c, d) to examine the behavior of specific pairs. All distances were smoothed with a 10ns rolling average.

### Sequence alignments

Multiple sequence alignment was performed with Clustal Omega^91^.

### Figures and movie preparation

All structure figures were produced with UCSF Chimera^92^, ChimeraX, VMD and PyMol (http://www.pymol.org). Pore radius calculations were performed with HOLE implemented in *Coot*^93^. Movies were created using Chimera^92^ and VMD^94^. Figures were composited in Adobe Photoshop and PowerPoint. Movies were composited in VideoMach.

## Data availability

Data supporting the findings of this study are available from the corresponding author upon reasonable request. A reporting summary for this Article is available as a Supplementary Information file. CryoEM density maps have been deposited to the Electron Microscopy Data Bank (P1: EMD-23864, P1: EMD-23881, Q1: EMD-23880 and Q2: EMD-23882). Coordinates for (Kir6.1)_4_-SUR2B atomic models have been deposited to the Protein Data Bank (P1: 7MIT, P2: 7MJP, Q1: 7MJO and Q2: 7MJQ).

## Acknowledgements

We thank Dr. Camden Driggers for helpful discussion. A portion of this research was supported by NIH grant U24GM129547 and performed at the Pacific Northwest Cryo-EM Center (PNCC) at Oregon Health and Science University and accessed through EMSL (grid.436923.9), a DOE Office of Science User Facility sponsored by the Office of Biological and Environmental Research. We also thank Dr. Nancy Meyer at the PNCC and staff at the Multiscale Microscopy Core (MMC) of Oregon Health & Science University (OHSU) for technical support. We acknowledge the support by the National Institutes of Health grant R01DK066485 (to S.-L. S.) and by the National Science Foundation grant MCB 17158233 (to D.M.Z.).

## Author contributions

MWS conducted the protein purification and reconstitution of CryoEM specimens, collected the CryoEM datasets, performed image analysis, atomic modeling, MD simulation analysis, and wrote the manuscript. ZY constructed the adenoviruses and performed protein expression and biochemical experiments. BLP wrote the manuscript. BM and JR conducted and analyzed the MD simulations. BM, JR and DMZ contributed to the experimental design and analysis of MD simulations. SLS conceived the project, provided overall guidance to the design and execution of the work, and wrote the manuscript. All authors contributed to manuscript preparation.

## Abbreviations

ABC: ATP-Binding Cassette
CD: Cytoplasmic Domain
CTD: C-Terminal Domain
Glib: Glibenclamide
KNt: Kir6.x N-terminal domain
NBD: Nucleotide Binding Domain
TM: TransMembrane
TMD: TransMembrane Domain
TMB: TransMembrane Bundle

---

**Supplementary Table 1.** Statistics of cryo-EM data collection, 3D reconstruction and model building.

|  |  |  |  |  |
| --- | --- | --- | --- | --- |
| Data collection |  |  |  |  |
| Microscope | Krios |  |  |  |
| Voltage (kV) | 300 |  |  |  |
| Camera | Gatan K3 |  |  |  |
| Camera mode | Super-resolution |  |  |  |
| Defocus range (μm) | -1.3 ~ -2.8 |  |  |  |
| Movies | 4130 |  |  |  |
| Frames/movie | 70 |  |  |  |
| Exposure time (s) | 7 |  |  |  |
| Dose rate (e<sup>-</sup>/pixel/s) | 16 |  |  |  |
| Magnified pixel size (Å) | 1.653* |  |  |  |
| Total Dose (e<sup>-</sup>/Å<sup>2</sup>) | ~41 |  |  |  |
| Reconstruction |  |  |  |  |
| (Kir6.1)4SUR2B focused refinement | P1 | P2 | Q1 | Q2 |
| Software | Relion 3.0 | Relion 3.0 | Relion 3.0 | Relion 3.0 |
| Symmetry | C1 | C1 | C1 | C1 |
| Particles refined | 139,944 | 70,830 | 71,880 | 22,038 |
| Resolution (masked) | 3.4.Å | 4.2Å | 4.0 Å | 4.2Å |
| Model Statistics |  |  |  |  |
|  | (full KNt) | (full KNt) | (short KNt)** | (short KNt)** |
| Map CC (masked) | 0.80 | 0.65 | 0.77 | 0.66 |
| Clash score | 8.08 | 2.43 | 8.25 | 5.92 |
| Molprobity score | 1.87 | 1.55 | 1.97 | 1.86 |
| Cβ deviations (%) | 0 | 0 | 0 | 0 |
| Rotamer (%) | 0 | 0 | 0 | 0 |
| Ramachandran |  |  |  |  |
| Outliers (%) | 0 | 0 | 0 | 0 |
| Allowed (%) | 6.62 | 9.11 | 8.99 | 9.43 |
| Favored (%) | 93.38 | 90.89 | 91.01 | 90.57 |
| RMS deviations |  |  |  |  |
| Bond length | 0.006 | 0.006 | 0.004 | 0.010 |
| Bond angles | 0.661 | 1.002 | 0.710 | 1.461 |

\*Super-resolution pixel size 0.86.
\*\*short KNt: Missing aa 1-21 for Q1 and 1-22 for Q2.

**Figure S1.**
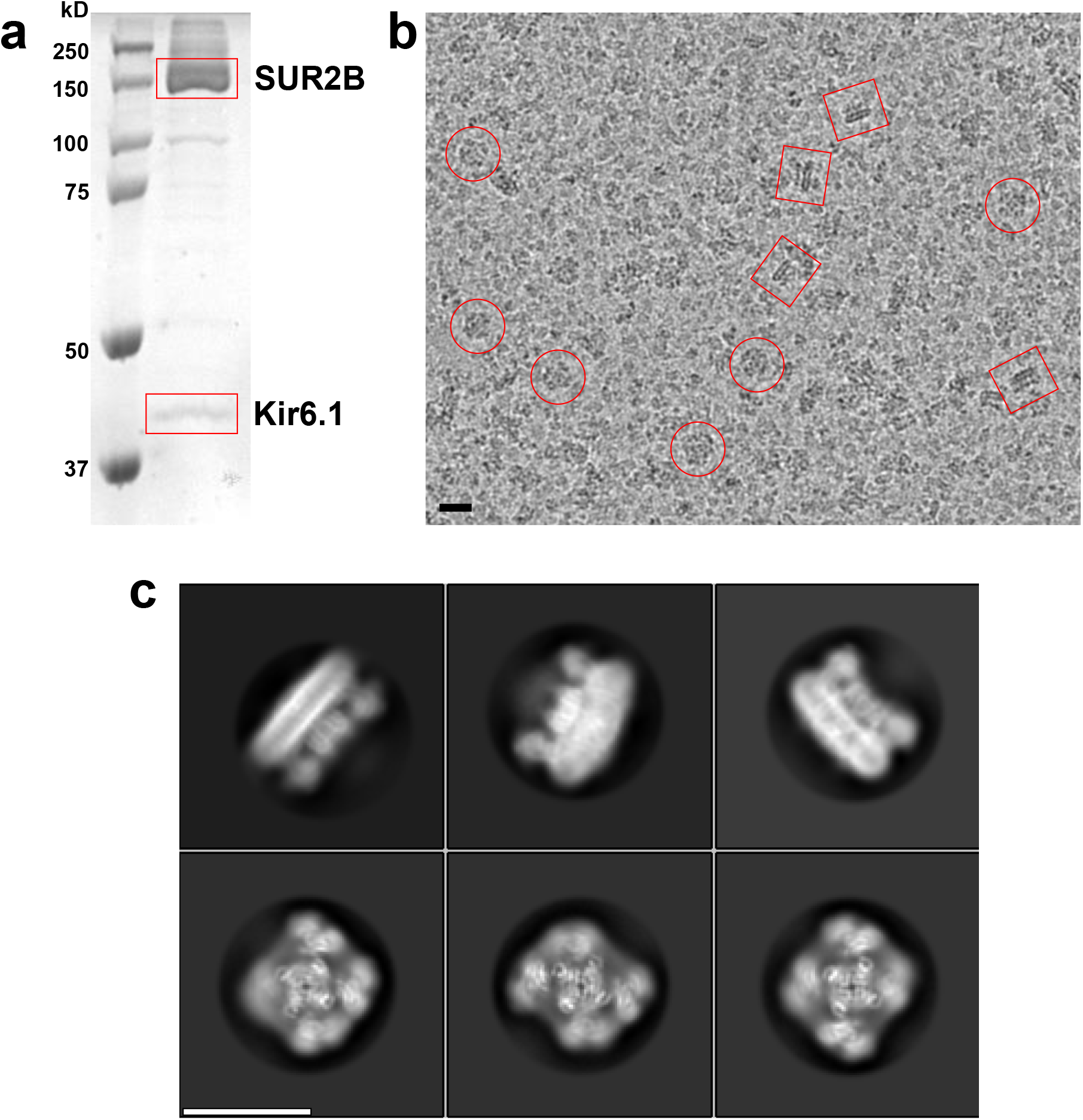
(a) Coomassie gel showing purified SUR2B and Kir6.1. Note SUR2B has two bands: the upper band corresponds to complex-glycosylated mature protein and the lower band the core-glycosylated immature protein. (b) CryoEM micrograph showing single K_ATP_ channel particle in top/down view (red circle) and side view (blue circle). (c) Examples of 2D class average. Scale bars:20nm.

**Figure S2.**
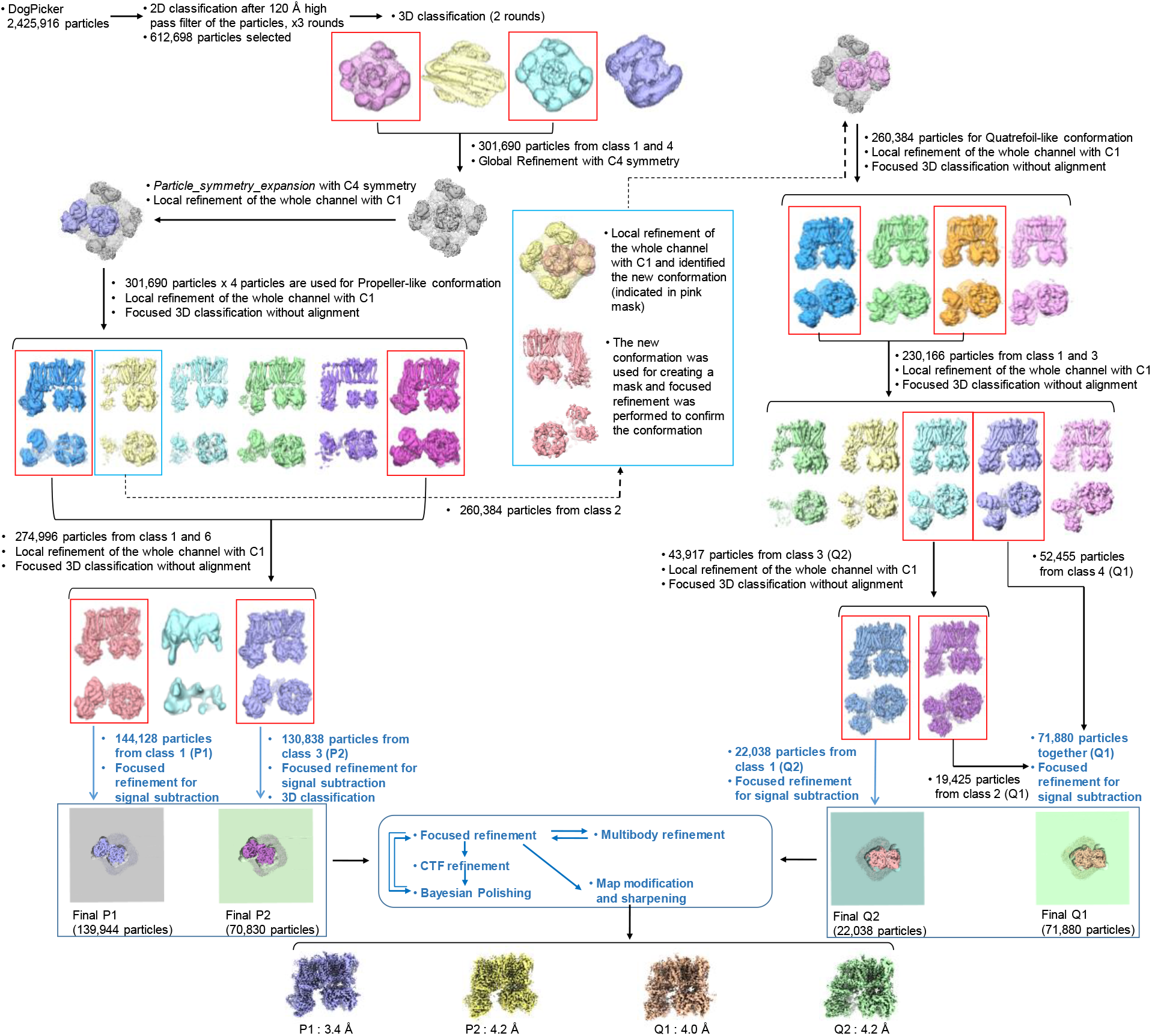
Image data processing workflow.

**Figure S3.**
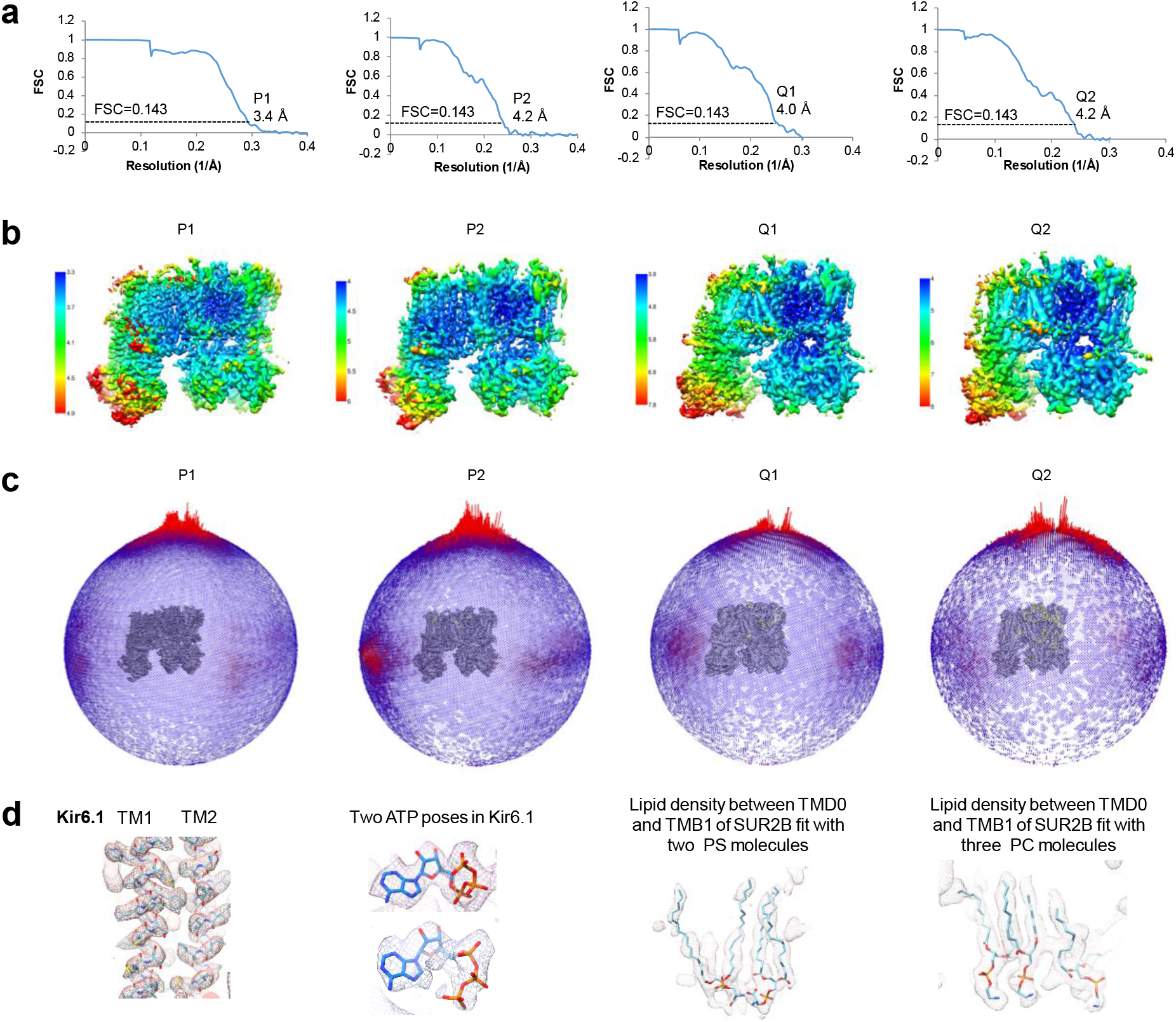
(a) FSC curves for the four conformation classes P1, P2, Q1, and Q2. (b) Local resolution for the four conformation classes. (c) Particle distributions of the four 3D classes. (d) Examples of cryoEM density fitting for protein, ligands, and lipids observed in the Kir6.1/SUR2B channel map.

**Figure S4.**
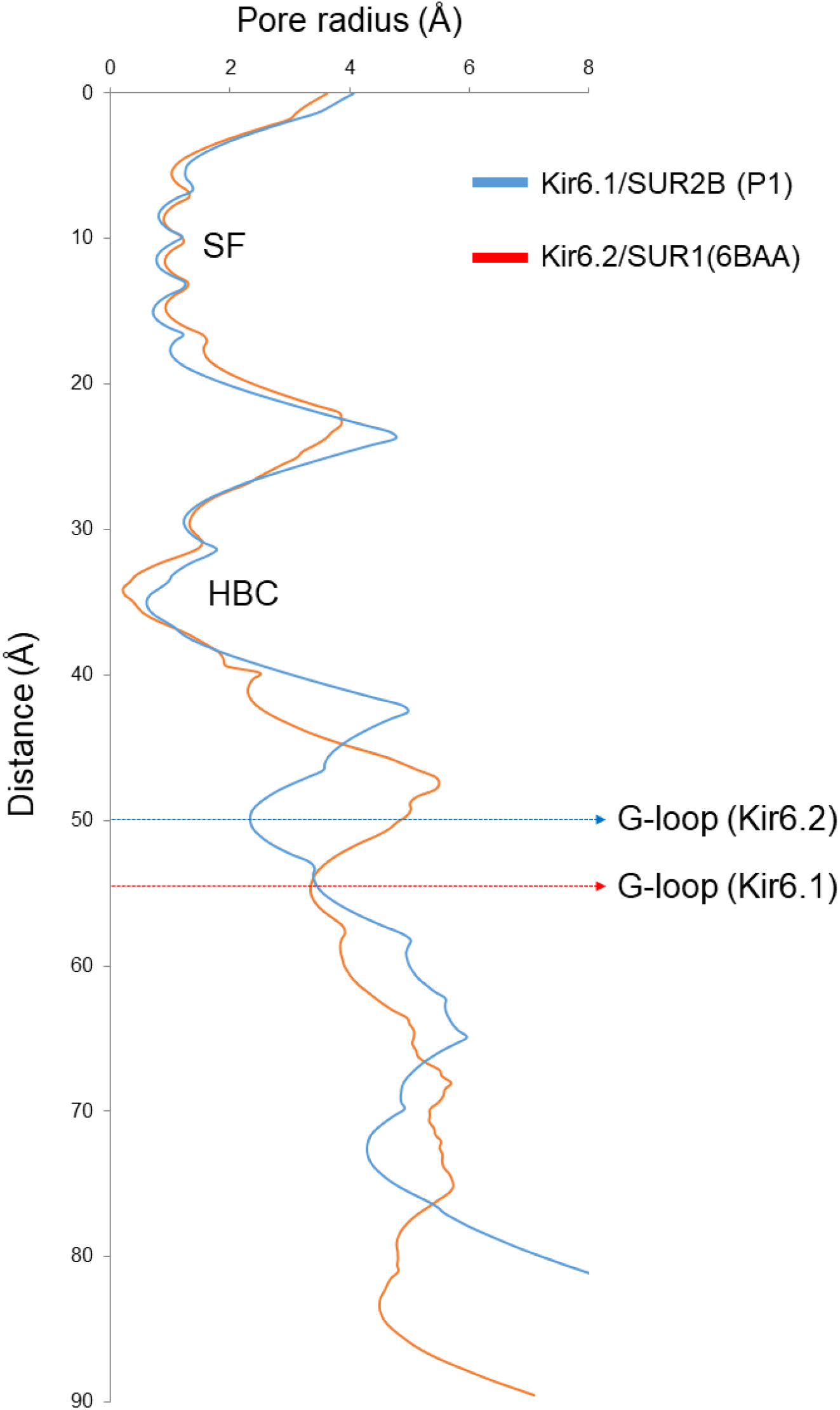
Pore radius plot for Kir6.1 (blue) and Kir6.2 (red). Note the G-loop gate is significantly further away from the HBC gate in Kir6.1 than in Kir6.2.

**Figure S5.**
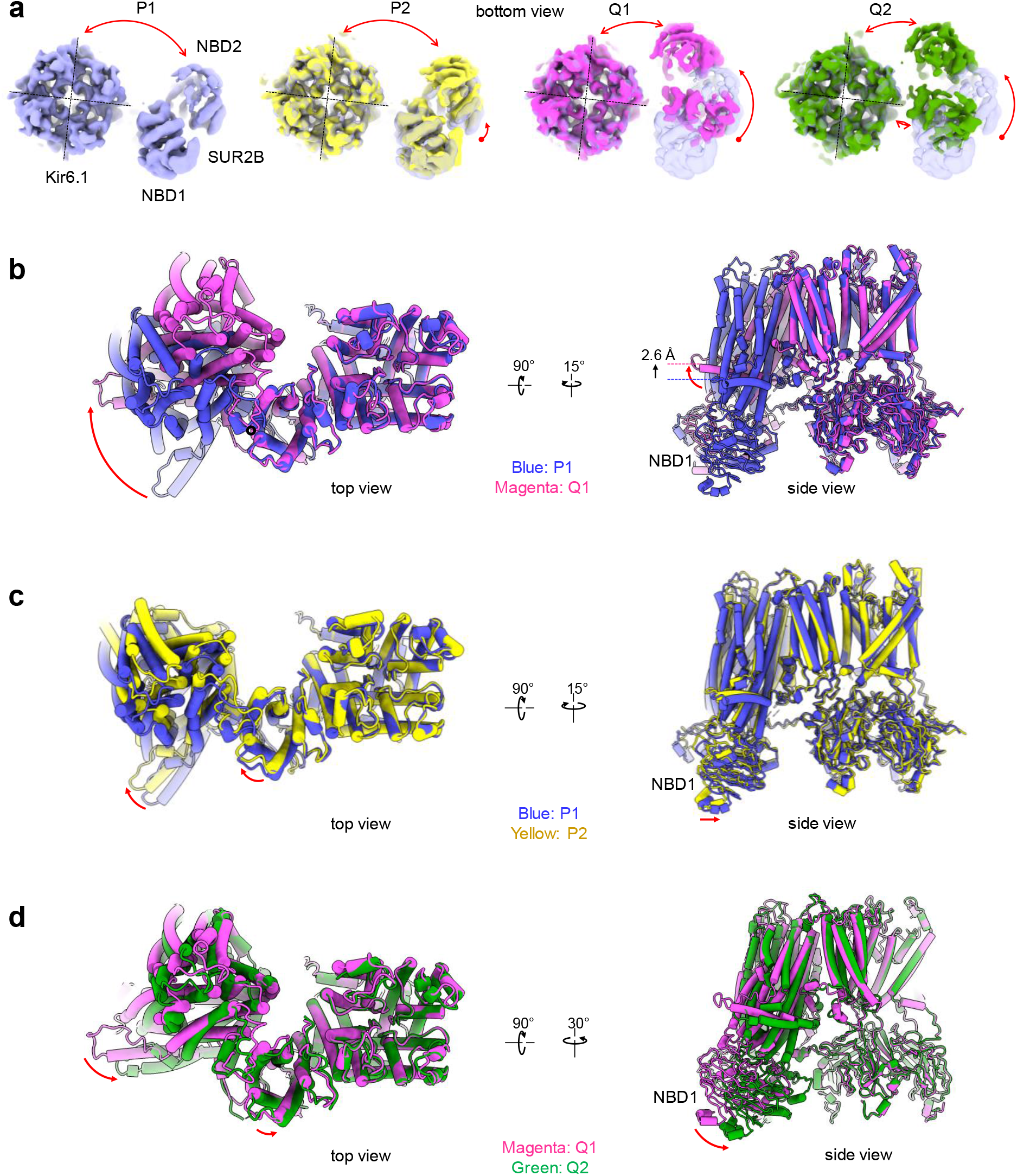
(a) Bottom view of the four different conformations showing the variation in rotation and distance of the two NBDs relative to the Kir6.1 tetramer (double red arrows). The one direction red arrows in P2, Q1, and Q2 indicate the difference in the NBDs relative to P1 (transparent blue). (b) Overlay of P1 and Q1 structures. (c) Overlay of P1 and P2 structures. (d) Overlay of Q1 and Q2 structures. The red arrows in b, c, and d mark the differences of the two superimposed structures.

**Figure S6.**
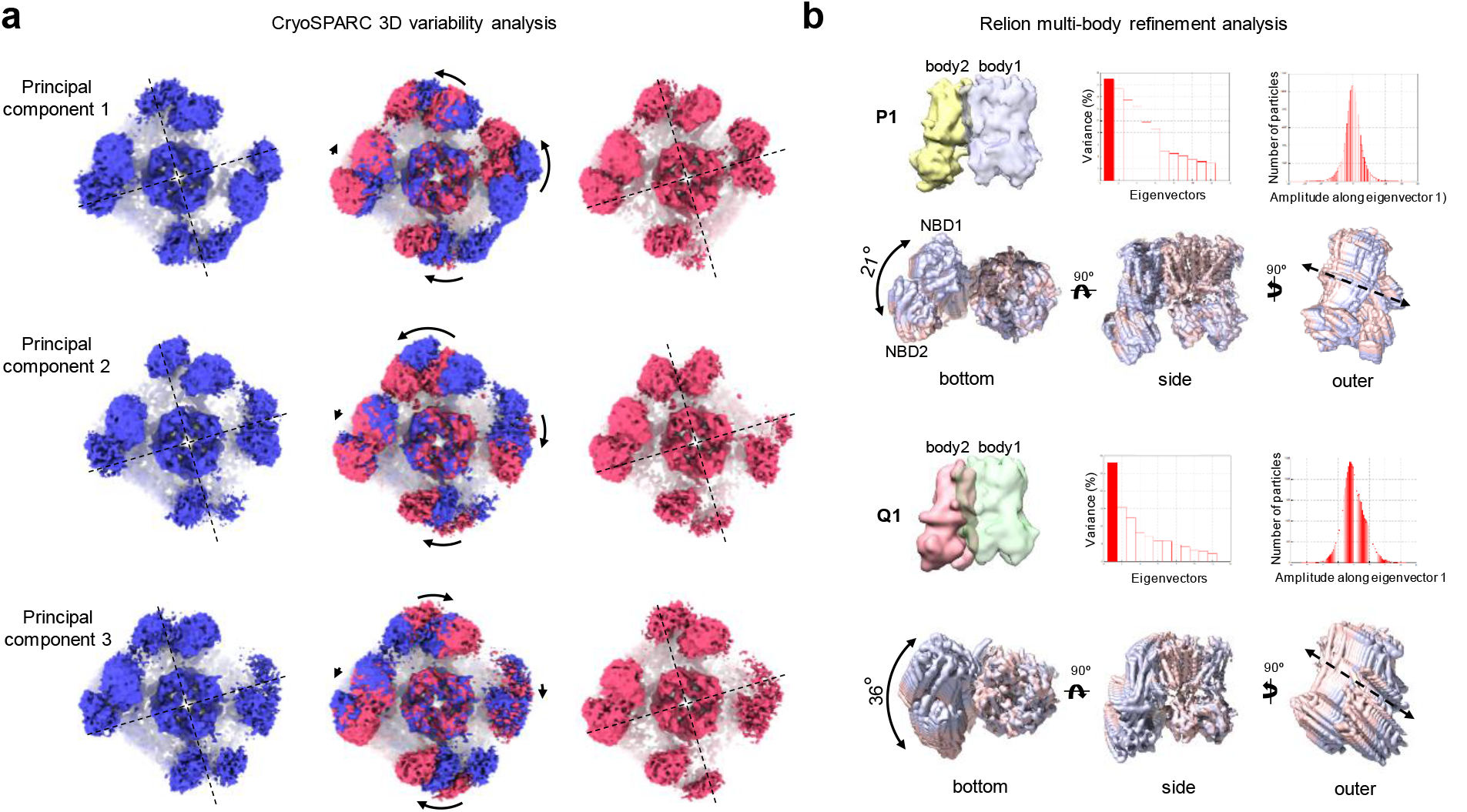
(a) CryoSPARC 3D variability analysis reveals dynamic movements of the SUR2B subunits. Shown are bottom view of conformational variability observed in three variability components (top: principal component 1; middle: principal component 2; bottom: principal component 3). Blue and red on the left and right columns represent the extreme ends of the conformation variation and the center column is the overlay of the two conformations, with arrows indicating directions of movement from blue to red. (b) Multibody refinement analysis showing heterogeneity of the SUR2B subunits in P1 and Q1 conformations. For each, assignment of the two bodies, plot of eigenvectors contributing to the variance observed, particle distribution plot along eigenvector 1 (red column in eigenvector plot) are shown on the top. The maps below show the dynamic range seen in each conformation along the first eigenvector, with light blue representing 50% of the particles from the left, and light pink representing 50% of the particles from the right side of the particle distribution plot.

**Figure S7.**
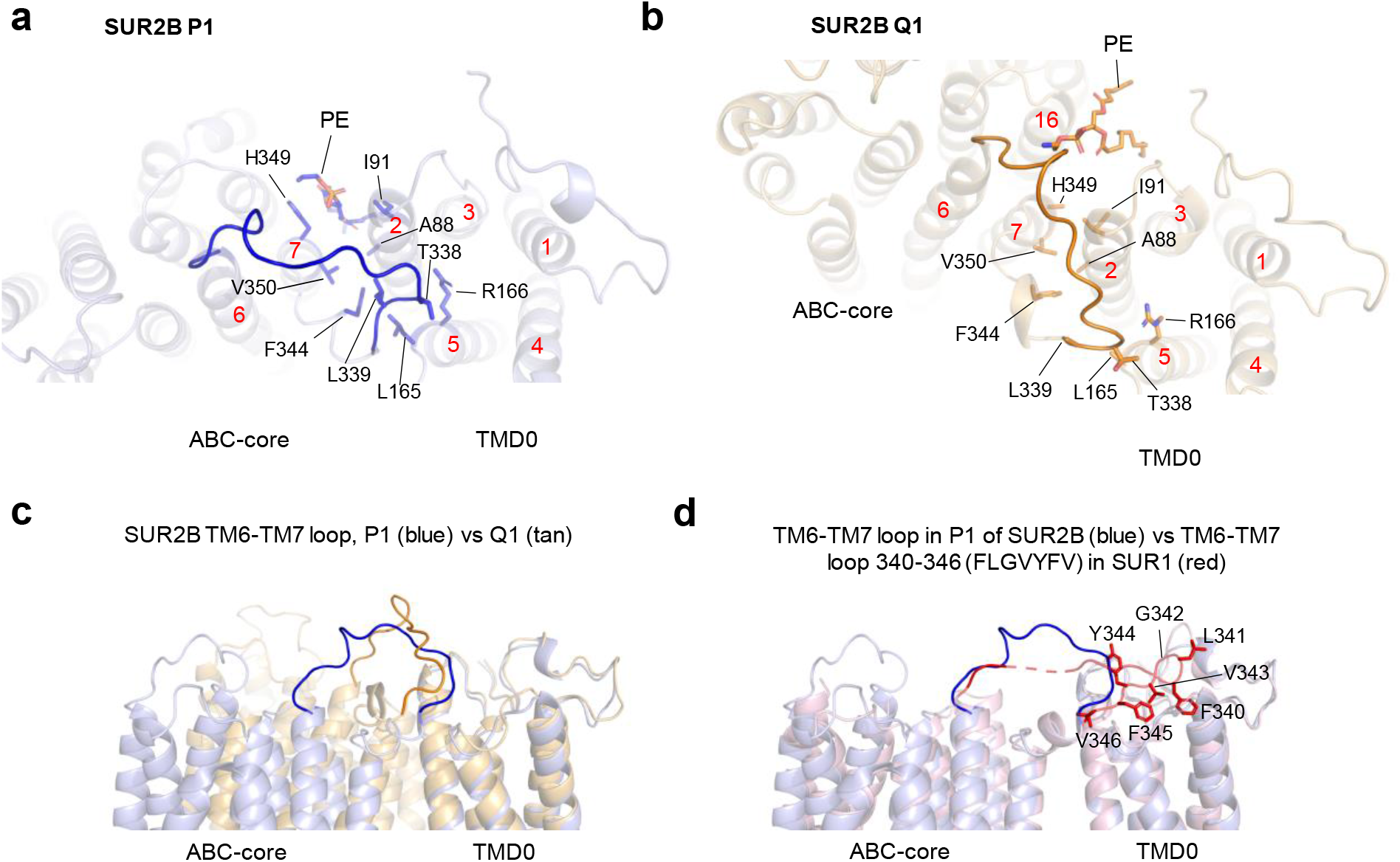
Interactions between the ABC core and TMD0 viewed from the extracellular side in P1 (a) and Q1(b) conformations. PE: PhosphatidylEthanolamine. The red numbers indicate the number of transmembrane helices. (c) Superposition of the P1 and Q1 structure and viewed from the side showing the difference in the loop between TM6 and TM7. (d) Superposition of SUR2B in P1 conformation and SUR1 viewed from the side. SUR1 contains a series of hydrophobic residues that interact with a hydrophobic pocket in TMD0. All models are aligned against the TMD of Kir6.1.

**Figure S8.**
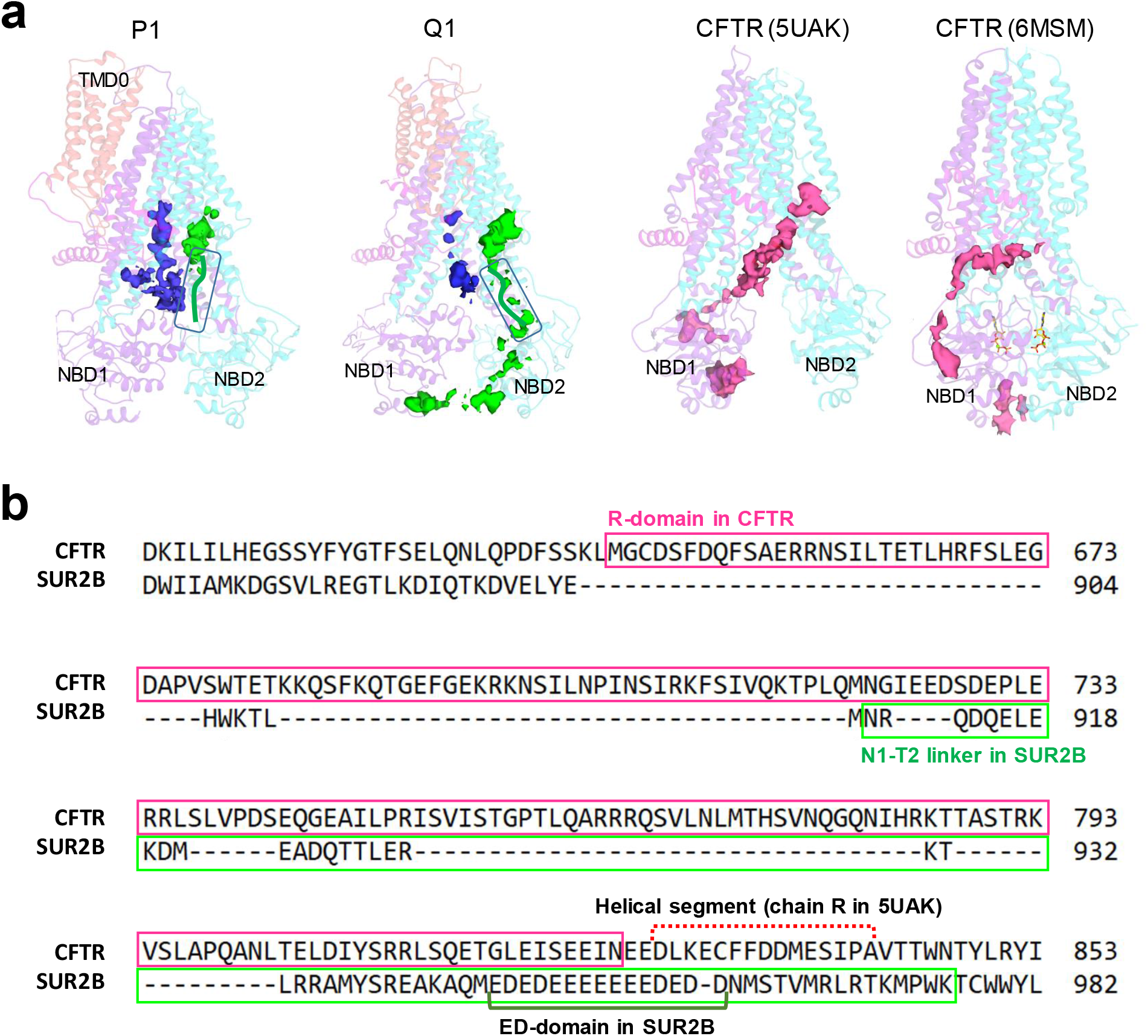
(a) Comparison of the cryoEM density of the N1-T2 linker in SUR2B P1, Q1 models (ED domain boxed in green) and corresponding linker in human CFTR with the R-domain unphosphorylated and NBDs separate (PDB: 5UAK, EMD-8516) or R-domain phosphorylated and NBDs dimerized (PDB:6MSM). The density of Kir6.1 N-terminus (blue) is also shown in SUR2B models as a reference. (b) Sequence alignment of CFTR R domain including a helical segment modeled in 5UAK, and N1-T2 linker of SUR2B including the ED-domain of 15 consecutive negatively charged glutamate and aspartate residues.

**Figure S9.**
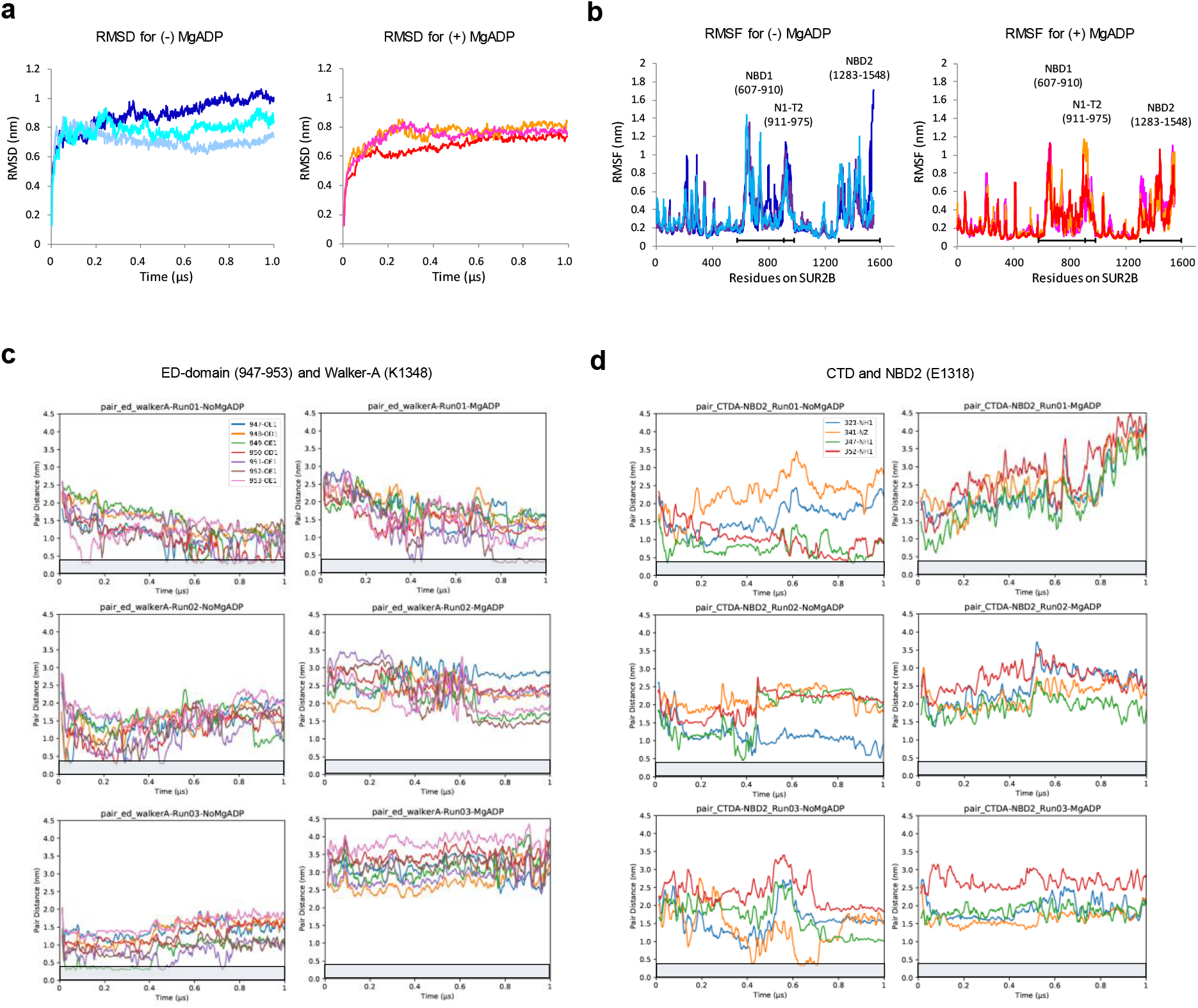
(a) RMSD of the three separate runs under each condition. (b) RMSFs of SUR2B residues in three 1μs simulation runs under the (-) MgADP or (+) MgADP condition (see Fig.S9a). Regions that are highly dynamic are marked below the traces with double arrows and labeled above the traces. (c) Measurement of distance between side chain oxygen in the glutamate/aspartate of the ED-domain residues 947-953 and side chain nitrogen of K1348 in the three individual runs under both conditions. (d) Same as (c) except the distance measured is between side-chain nitrogen of Kir6.1-CTD residues R323, K341, R347, R352, and the side chain oxygen of E1318. The grey box in each plot in (c) and (d) marks the region with distance ≤ 4Å.

**Video 1**. Morph of Kir6.1-tetramer + SUR2B structure in P1, P2, Q1 and Q2 conformations. The models in P1, P2, Q1 and Q2 are aligned against TMD of Kir6.1 (60-185) and only protein parts are shown with the same color scheme as in Figure 1.

**Video 2**. 3D variability of whole channel viewed from the cytoplasmic side showing asynchronous dynamics of the ABC-core of the four SUR2B subunits. The map was low-pass filtered to 5 Å.

**Video 3**. Morph videos of eigen vector 1 from Multibody refinement of P1 and Q1 conformations showing relative movement of body 1 (Kir6.1 tetramer + TMD0) and body 2 (ABC-core of SUR2B).

**Video 4**. Morph video showing local structural changes in L0 loop in P1, P2, Q1 and Q2 conformations. Lipids in P1 conformation disappear when part of L0 loop (197-224) moves up towards the membrane in Q1 conformation. Alignment of the models are the same as Video1.

**Video 5**. Representative MD simulations (run 1) of Kir6.1 tetramer + SUR2B in the absence of MgADP at NBD2. The start model is aligned against TMD of Kir6.1 (60-185) throughout the trajectory with a frame rate 200/μs. The trajectory is smoothed with step 1.

**Video 6**. Representative MD simulations (run 2) of Kir6.1 tetramer + SUR2B in the presence of MgADP at NBD2 at the start of the simulation. The start model is aligned against TMD of Kir6.1 (60-185) throughout the trajectory with a frame rate of 200/μs. The trajectory is smoothed with step 1.

